# Gonadal Hormone Status Dictates Neurobehavioral, Metabolic and Immunological Benefits of Ketogenic Diet in Female Mice through Gut-Metabolic-Brain Axis

**DOI:** 10.1101/2025.11.28.691132

**Authors:** Anil Kumar Rana, Nisha Gautam, Md Minhajul Abedin, Parampal Singh, Amit Kumar Rai, Jyotdeep Kaur, Mohit Kumar

## Abstract

The ketogenic diet (KD) is recognized for its broad therapeutic potential across metabolic, neurological, immunological, and aging-related domains; however, its efficacy and durability in females remain underexplored. This study investigated the effects of a short-term, intermittent KD on neurobehavioral function, lipid metabolism, immune response, and gut microbiota composition in sham-operated (intact) and ovariectomized (OVX, estrogen-deficient) female mice, aiming to delineate the interactions between diet and gonadal hormones. In sham females, KD reduced circulating estradiol levels and induced low-grade systemic inflammation, without producing measurable behavioral benefits. Conversely, KD exhibited reduced compulsivity, anxiety, and depression-like behaviors with improved well-being in OVX females, suggesting enhanced neurobehavioral resilience under estrogen-deficient conditions. Biochemical analyses revealed KD-induced hyperlipidemia in both groups, although the lipid load was attenuated in OVX mice. KD further decreased low-density lipoprotein cholesterol but led to uremia and histopathological evidence of hepatic steatosis and fibrosis in OVX females, despite unaltered liver function markers. 16S rRNA sequencing revealed hormone-dependent remodelling of the gut microbiota, characterised by distinct compositional shifts between the sham and OVX cohorts. Correlation analyses linked microbial alterations to improved lipid profiles and reduced anxiodepressive behaviors, implicating the gut-metabolic-brain axis in mediating the effects of KD. Collectively, these findings suggest that KD confers neurobehavioral and metabolic benefits in the absence of ovarian hormones but may also pose risks to the liver and kidneys. These results highlight ketogenic nutrition as a potential non-pharmacological strategy to mitigate menopause- and aging-associated neurobehavioral, metabolic, and immune dysfunctions through modulation of the gut microbiota.

## Introduction

A ketogenic diet (KD), characterized by high fat, adequate protein, and low carbohydrate intake, induces a metabolic state of ketosis, where the body primarily utilizes ketone bodies for energy in the absence of sufficient glucose [1]. This metabolic shift has been linked to improved glucose and lipid metabolism, reduced triglycerides and total cholesterol, elevated high-density lipoprotein levels, and enhanced immune functions in both health and disease [2–5]. Initially established as an effective intervention for intractable epilepsy, the KD has gained increasing attention for its potential therapeutic roles in metabolic diseases such as obesity and type 2 diabetes [6,7], neurodegenerative diseases such as Alzheimer’s disease [8,9] and Parkinson’s disease [10,11] and certain psychiatric conditions such as Schizophrenia, Bipolar and major depressive disorder [12,13]. Given its broad range of beneficial effects, the KD is being actively investigated as a practical therapeutic intervention in multiple health domains.

Emerging evidence indicates that biological sex is a critical confounding factor shaping adaptive and therapeutic responses to KD, with females frequently exhibiting attenuated or distinct metabolic and physiological outcomes than males [14,15]. For example, a preclinical study has shown a pivotal role of gonadal hormones in modulating adaptive responses to KD [14], where male mice on KD exhibited significant weight loss and improved insulin sensitivity, but orchidectomy abolished these benefits, indicating that the effects were dependent on testicular hormones. In contrast, female mice on KD gained more weight and exhibited no improvement in insulin sensitivity compared to controls. Interestingly, ovariectomy reversed this trend, as females lost weight on KD but remained glucose intolerant. Furthermore, KD induces low-grade inflammation and cellular senescence across multiple organs in male mice, driven by increased oxidative stress through acetylation of the mitochondrial superoxide dismutase enzyme. These detrimental effects were mitigated in estradiol-supplemented males and completely prevented in females, underscoring a sex-specific adaptive response to KD and highlighting the critical role of gonadal hormones in shaping its metabolic outcomes [15]. Similarly, a recent study shows a male-specific anti-inflammatory effect of KD in a mouse model of allergic airway inflammation [16]. Our previous studies also show that KD did not induce behavioral and neuroimmune changes in ovary-intact female mice that were evident in age-matched male counterparts [17], raising the fundamental question of whether female gonadal hormones, primarily estrogen, regulate their adaptive response to KD among different health domains.

Current literature provides only fragmented insights, often focusing on individual organ systems or relying on single-sex models, thereby limiting a comprehensive view of the systemic effects of KD in females. There is a growing need to determine how the absence of gonadal hormones alters the risks and benefits of KD in females, a question central to developing effective metabolic therapies for women’s health.

## Results

### KD suppresses systemic estradiol levels in sham females

In our recent preclinical study, we found that the beneficial effects of the KD on behavioral outcomes, particularly anxiety- and depressive-like behaviors, as well as on neuroinflammatory responses, were sex-specific.

Specifically, KD enhanced behavioral vitality and reduced neuroinflammation in male mice, but not in females [17]. In this study, we sought to determine whether depletion of female gonadal hormone, especially estrogen, could modulate the adaptability of KD in females. For that, female mice had either bilateral OVX surgery, in which both of their ovaries were surgically removed, or sham surgery, in which an abdominal incision was made and closed without the ovaries being removed. After two months of surgery, the animals were randomly assigned to either the CD or KD for the next four weeks, as shown in **Fig. 1A**.

**Figure 1:**
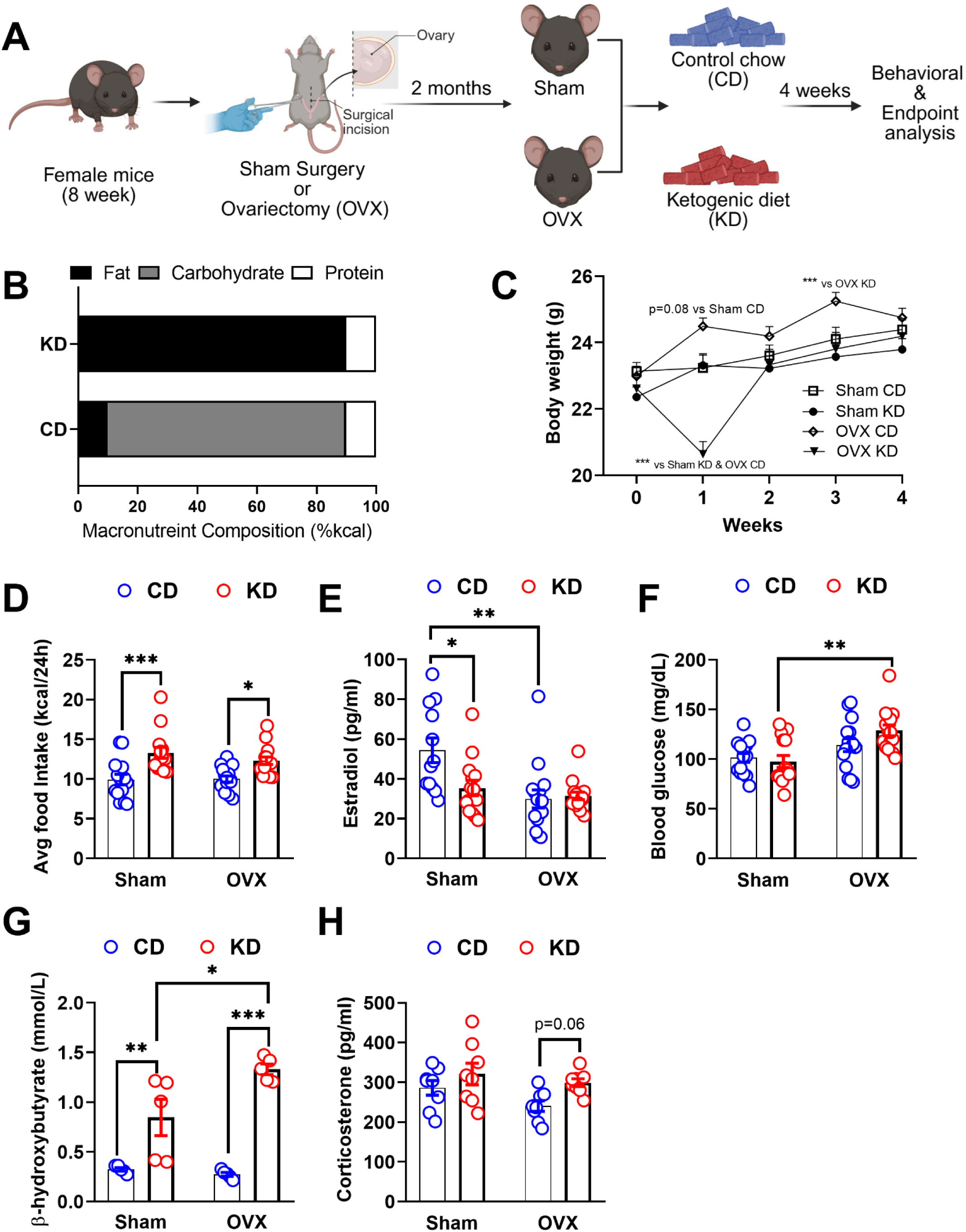
KD reduces estradiol levels in sham female mice while inducing blood glucose levels in OVX females. **(A)** Schematic representation of the study timeline. **(B)** Percentage of macronutrients in the CD and KD. (**C**) Body weight of animals throughout the study. **(D)** Average food intake (kcal/24h). **(E)** Plasma estradiol levels. **(F)** Fasting blood glucose levels. **(G)** Plasma β-hydroxybutyrate levels. **(H)** Plasma corticosterone levels after four weeks of CD and KD in sham and OVX female mice. All data are expressed as mean ± SEM. Statistical significance is indicated as *p<0.05, ** p<0.01, *** p<0.001, and **** p<0.0001. Sample size: Panel C-F, n = 14 for Sham CD and Sham KD; n = 15 for OVX CD and OVX KD. Panel G, n = 5 per group. Panel H, n = 8 per group. **CD**: Control diet; **KD**: Ketogenic diet; **OVX**: Ovariectomy; **Sham CD**: Group subjected to the sham surgery with a control diet; **Sham KD**: Group subjected to the sham surgery with a ketogenic diet; **OVX CD**: Group subjected to the OVX surgery with a control diet; and **OVX KD**: Group subjected to the OVX surgery with a ketogenic diet.

KD did not affect the body weight of sham females; however, a transient drop in body weight was observed in OVX females after one week of KD as compared to their OVX CD counterparts **(Fig.1C)**. Though those females regained the body weight by the second week, which then chronologically increased, aligning with the weight trajectories of the other experimental groups throughout the study. Both sham and OVX female mice on KD consumed more calories as compared to their CD counterparts **(Fig. 1D)**.

Serum estradiol levels were measured to assess the impact of KD on ovarian hormones in female mice **(Fig. 1E)**. As expected, OVX resulted in a drastic decline in blood estradiol levels as compared to sham controls. Surprisingly, KD also reduced blood estradiol levels in the sham females compared to their CD counterparts, suggesting that a brief intermittent KD may either disrupt the hypothalamic-pituitary-ovarian axis, inhibit estrogen synthesis or compromise overall estrogen metabolism in the surrounding tissues of ovaries-intact females.

KD did not affect fasting blood glucose levels in sham and OVX female mice; however, blood glucose levels were higher in OVX females on KD as compared to their sham KD counterparts **(Fig. 1F)**. Elevated levels of plasma BHB in KD-fed sham and OVX females confirmed the nutritional ketosis induced by KD. However, OVX females on KD had higher BHB levels as compared to their sham KD counterparts **(Fig. 1G)**. These findings suggest that four weeks of intermittent KD effectively induces nutritional ketosis in ovaries-intact as well as OVX females and that KD induces hyperglycaemia and further increases blood ketone levels in ovarian hormone-deprived females.

Serum corticosterone levels were also examined to check the adaptability of KD in females **(Fig. 1H)**. No change in systemic corticosterone levels was found in sham and OVX females after transitioning to KD.

### KD increases the behavioral vitality in OVX female mice

Following four weeks of KD, OFT and EPM were used to assess locomotion and anxiety-like behavior in all treatment groups. During OFT, KD did not affect the amount of time spent in the central zone in sham females. However, OVX females on KD spent more time in the center zone during OFT compared to their CD counterparts, suggesting anxiolytic effects of KD in OVX female mice **(Fig. 2A)**. The time spent in the peripheral area during the OFT is another measure of anxiety-like behavior in rodents, and it is directly linked to anxiety levels. OVX females on KD spent less time in the periphery during OFT than their CD and sham KD counterparts **(Fig. 2B)**. However, no difference in time spent in the peripheral zone was found between sham females fed either CD or KD. KD did not improve locomotion in either sham or OVX females; however, sham females on KD traveled a greater distance than their OVX KD counterparts **(Fig. 2C)**, suggesting that the effect of KD on spontaneous locomotor activity is dependent on the status of ovarian hormones in females.

**Figure 2:**
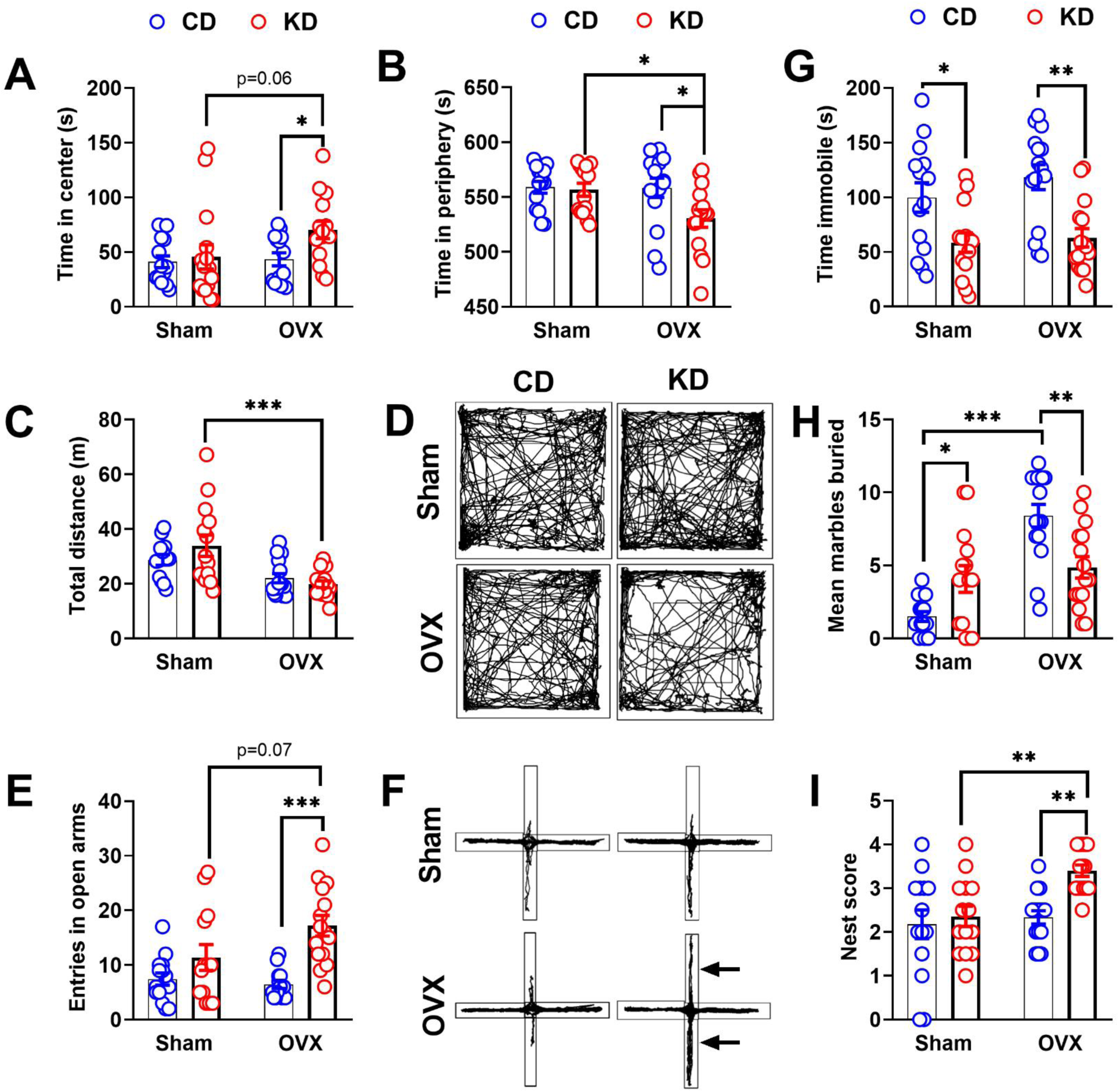
Effects of KD on neurobehavioral outcomes in sham and OVX female mice. (**A-D) OFT:** Analysis of general locomotor activity and anxiety-like behavior. **(A)** Time spent in the center zone (s), **(B)** Total time spent in the peripheral zone (s), **(C)** Total distance traveled (m), and (**D**) Representative tracks plots. (**E-F) EPM:** Assessment of anxiety-like behavior. **(E)** Number of entries into the open arms and (**F**) Representative track plots. **(G) FST:** Total immobility time (s), a measure of depression-like behavior. **(H) MBT:** Number of marbles buried, a measure of repetitive or compulsive-like behavior. (**I**) **Nesting Behavior**: Total nest score, reflecting an animal’s ability to build a complex nest, often indicative of well-being, cognitive function, and motivated behavior. Scores are typically graded on a scale (e.g., 0-5), with higher scores indicating better nesting. All data are expressed as mean ± SEM. Statistical significance is indicated as *p<0.05, ** p<0.01, *** p<0.001, and **** p<0.0001. Sample size: Panel A-B, n =14 for Sham CD and Sham KD; n = 15 for OVX CD and OVX KD. Panel C, n = 13 for Sham CD; n = 14 for Sham KD; and n = 15 for OVX CD and OVX KD. Panel E, n = 14 for Sham CD; n = 13 for Sham KD and OVX CD; and n = 15 for OVX KD. Panel G-I, n = 14 for Sham CD and Sham KD; n = 15 for OVX CD and OVX KD. **CD**: Control diet; **KD**: Ketogenic diet; **OVX**: Ovariectomy; **Sham CD**: Group subjected to the sham surgery with a control diet; **Sham KD**: Group subjected to the sham surgery with a ketogenic diet; **OVX CD**: Group subjected to the OVX surgery with a control diet; **OVX KD**: Group subjected to the OVX surgery with a ketogenic diet; **MBT**: Marble burning test; **OFT**: Open field test; **EPM**: Elevated plus maze; **FST**: Forced swimming test; and **s** = seconds.

To confirm the behavioral data from the OFT, we performed an EPM test to assess anxiety-like behavior **(Fig. 2E)**. The number of entries in open arms increased significantly in OVX females on KD when compared to CD counterparts, indicating anxiolytic effects of KD in OVX females. Overall, OFT and EPM results show that KD decreases anxiety levels in OVX females but not in ovaries-intact females.

In contrast, the KD significantly improved depression-like behavior in both sham and OVX female groups, as measured by the FST **(Fig. 2G)**. This was demonstrated by a considerable decrease in immobility time among those females as compared to their CD counterparts. These results indicate that the KD had a primary effect on reducing depression-like behavior in female mice, regardless of their gonadal hormonal state.

Further, MBT was performed to evaluate the effect of KD on compulsive-like behavior. Sham KD and OVX CD females buried considerably more marbles than the sham CD females **(Fig. 2H)**, indicating the development of compulsive-like behavior in those animals. However, KD reduced the number of buried marbles in OVX females when compared to their CD counterparts, suggesting the anti-compulsive effect of KD in those females.

We also looked at how a KD affected nesting behavior in female mice as a measure of overall mental well-being. However, KD did not increase nest construction scores in sham females. OVX females given KD showed a significant improvement in nest formation score as compared to CD controls **(Fig. 2I)**. The behavioral findings indicate that a KD increases overall mental well-being primarily in OVX females.

### Transcriptomic remodeling of the medial prefrontal cortex (mPFC) by KD in OVX females

Based on the observed behavioral improvements by KD in OVX females, we investigated transcriptomic changes in the mPFC, a key region regulating mood and executive functions [18], by KD in OVX females. Bulk RNA-seq data revealed that KD caused a comprehensive remodeling of the mPFC transcriptome in OVX mice, including the differential expression of genes involved in synaptic signaling, neuroplasticity, neuroinflammation, epigenetics, and energy metabolism.

The altered gene expression profile was visualized in a comprehensive heatmap generated from gene ontology (GO)-enriched differentially expressed genes (DEGs) **(Fig. 3B)**. This analysis revealed a distinct transcriptional signature between OVX CD and OVX KD groups. OVX CD samples clustered tightly, characterized by widespread downregulation of a large subset of genes. In contrast, OVX KD samples formed a distinct cluster, with consistent upregulation of the same gene subset. This dichotomy illustrates a strong and reproducible effect of KD under OVX conditions.

**Figure 3.**
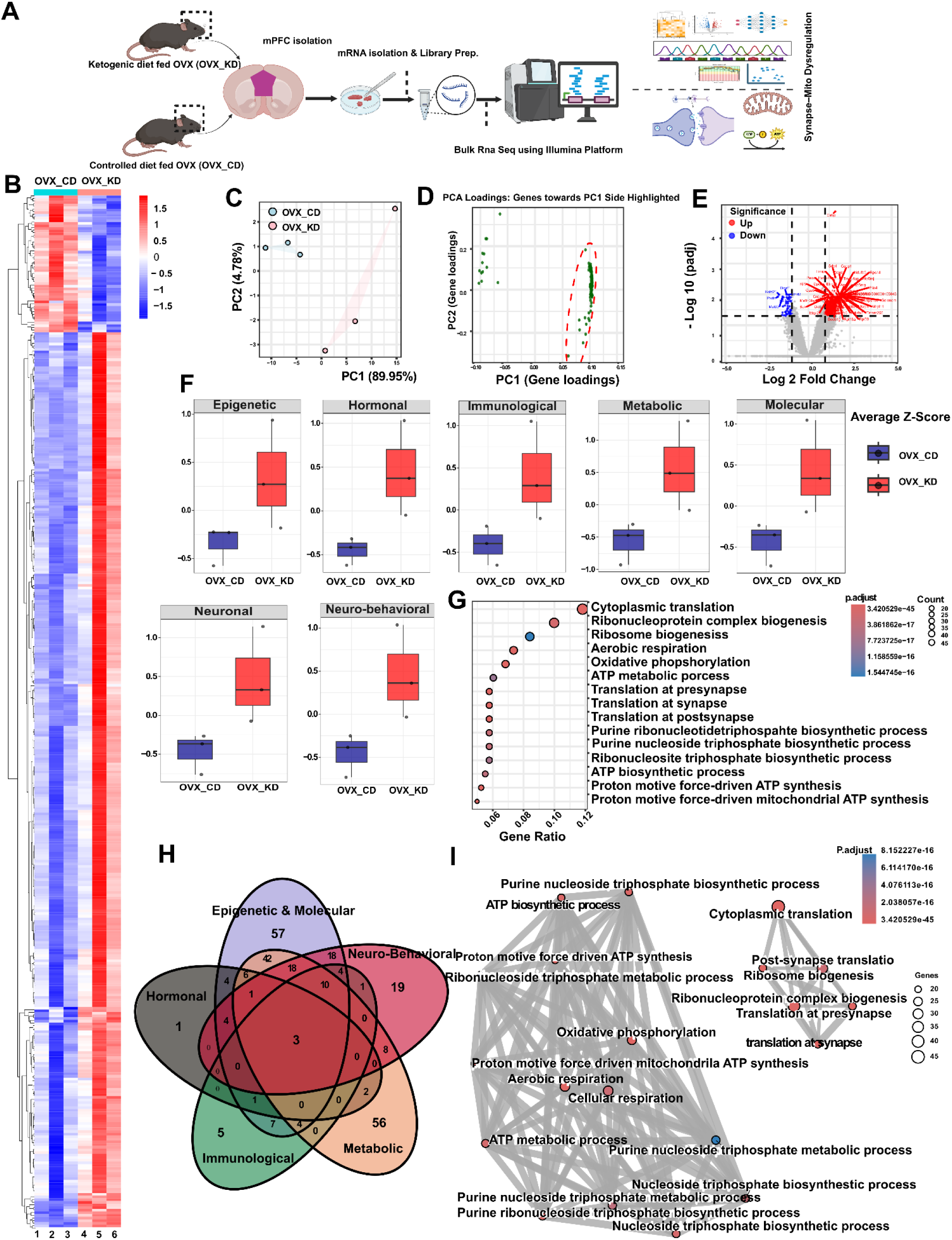
KD induces transcriptomic remodelling of the medial prefrontal cortex (mPFC) of OVX female mice. **(A)** Schematic overview of the transcriptomic analysis workflow. **(B)** Heatmap of GO-enriched DEGs (≈400). **(C)** PCA of normalized gene counts showing distinct separation of OVX KD (pink) and OVX CD (cyan) samples, with PC1 (≈89.9%) capturing diet-driven variance. **(D)** PCA loading plot highlighting DEGs contributing most to PC1, predominantly with positive loadings. **(E)** Volcano plot showing upregulated (red) and downregulated (blue) genes between OVX KD and OVX CD groups (|log₂FC| ≥ 1, padj < 0.05). **(F)** Boxplots of GO-module mean Z-scores indicating elevated activation in OVX KD group across epigenetic, metabolic, immunological, molecular, behavioral and neuronal pathways. **(G)** GO enrichment dot plot demonstrating upregulated processes such as translation, ribosome biogenesis, oxidative phosphorylation, and respiration. **(H)** Venn–ellipse plot showing overlapping genes across GO modules, with the strongest intersection between metabolic and epigenetic–molecular categories. **(I)** The GO network map illustrating functional connectivity between translation and metabolic pathways, revealing coordinated neuronal–metabolic adaptation in OVX KD mice.

To examine global transcriptomic variation and determine whether samples clustered according to biological groups, we performed principal component analysis (PCA) on normalized gene expression data **(Fig. 3C)**. This analysis revealed clear clustering of OVX KD samples separate from OVX CD, indicating a robust diet-driven transcriptomic shift under ovarian hormone deprivation. OVX KD replicates showed slightly greater dispersion within the group, possibly reflecting heterogeneity in biological response to the combined effects of KD and OVX. Despite this intra-group variation, the observed transcriptional changes remained consistent, confirming the robustness of downstream findings.

To quantify the extent of biologically relevant variation, we performed PCA focused specifically on the DEG set **(Fig. 3D)**. This analysis revealed a distinct clustering of genes along PC1, representing a high-confidence set of transcriptional changes driven by the interaction of diet and surgery. The convergence of heatmap, pathway enrichment, and DEG PCA analyses validates that the transcriptional alterations in the mPFC of OVX females on KD are both statistically and biologically significant and relevant.

To characterize specific transcriptional changes in the mPFC of OVX females by KD, we conducted differential gene expression analysis using DESeq2. The volcano plot **(Figure 3E)** displays log₂ fold change versus –log₁₀ adjusted p-values, highlighting significantly upregulated (red) and downregulated (blue) genes (padj; 0.05, |log₂FC| ≥1). A clear asymmetry in gene distribution was observed, with a greater number of upregulated transcripts in the OVX KD group. Prominent upregulated genes included *Txnip*, *Rpl3, Rps6, Ndufa10, Cox7a2l, Eif3b*, and *Mrpl15,* associated with translational and mitochondrial activity. In contrast, *Pfkfb3, Klf4* and *Pou3f3* were among the few downregulated genes, suggesting selective suppression of certain metabolic and transcriptional regulators. This pattern supports a pronounced anabolic and energy-augmenting transcriptional program in the mPFC of OVX females on KD.

Building on the PCA-based DEG analysis, we constructed GO modules by mapping DEGs to specific GO terms to define biologically meaningful categories. This analysis dissected the transcriptional changes into neuronal, behavioral, metabolic, epigenetic, molecular, hormonal and immunological modules. Module-specific heatmaps **(supplementary Fig. S1A)** and average gene expression Z-score **(Fig. 3F)** revealed clear transcriptional divergence between CD and KD-fed OVX females across nearly all domains, with KD exhibiting consistent upregulation of genes.

GO pathway enrichment analysis **(supplementary Fig. S1B)** further linked these transcriptional differences to functional changes. Notably, the mPFC of OVX females on KD exhibited a highly significant and consistent upregulation of pathways related to cytoplasmic translation, translation at presynapse, synapse, and postsynapse, indicating enhanced protein synthesis activity, particularly in neuronal compartments. Additional upregulated processes included oxidative phosphorylation, aerobic respiration, and proton motive force-driven mitochondrial ATP synthesis, suggesting increased mitochondrial bioenergetics potentially as a compensatory mechanism to meet elevated energy demands from enhanced translation and synaptic activity. Ribonucleoprotein complex biogenesis, ribosome biogenesis, and ATP biosynthesis were also enriched, supporting the notion of heightened anabolic processes. A smaller set of downregulated GO terms included negative regulation of DNA-binding transcription factor activity, negative regulation of G1/S transition of the cell cycle, and protein homo-oligomerization. The generation of GO pathway enrichment dot plots from the GO modules recapitulated earlier findings, confirming robust enrichment for pathways associated with synaptic regulation, translation, oxidative phosphorylation, aerobic respiration, metabolic processes, and ribonucleoprotein complex biogenesis **(Fig. 3G)**. Collectively, these pathway enrichments underscore a strong transcriptional reprogramming toward enhanced neuronal and metabolic activity in the mPFC of OVX females on KD.

Venn–ellipse analysis of GO modules revealed substantial intermodule overlap, indicating coordinated transcriptional regulation **(Fig. 3H)**. The epigenetic–molecular module contained 57 distinct genes, sharing significant overlap with metabolic and neuro-behavioral modules. The neuro-behavioral module contained 19 distinct genes, largely overlapping with epigenetic and metabolic modules. The metabolic module contained 56 distinct genes with substantial sharing across modules, highlighting the interconnected regulation of the mPFC transcriptome in OVX females on KD. Immunological and hormonal modules contained fewer distinct genes, but their presence further reflects the multi-dimensional impact of KD on the mPFC of OVX females.

Network analysis revealed tight interconnections between synaptic translation regulation and metabolic processes, including purine ribonucleotide biosynthesis and oxidative phosphorylation **(Fig. 3I)**. These results illustrate a coordinated functional architecture wherein neuronal and metabolic adaptations converge to drive the transcriptional reprogramming induced by KD in OVX mice. To probe these interconnections further, we performed cnetplot analysis **(supplementary Fig. S1D)**, which revealed the specific gene subsets driving these network-level enrichments. This visualization demonstrated that multiple DEGs co-map across synaptic and metabolic pathways, reinforcing their integrative regulatory roles and highlighting key transcriptional nodes that bridge neuronal and bioenergetic adaptations to KD in OVX females. These findings suggest that the KD-induced neurobehavioral changes observed in OVX females may be driven, at least in part, by epigenetic and metabolic alterations within the mPFC.

### KD induces hyperlipidaemia in female mice regardless of gonadal hormone status while reducing LDL levels in OVX females

As KD is a high-fat diet, we further examined the effect of KD on plasma lipid profile and liver health of sham and OVX female mice **(Fig. 4)**. Triglyceride levels were significantly higher in both sham and OVX females on KD compared to their CD counterparts **(Fig. 4A)**. Notably, a direct comparison of diet-matched groups found that OVX females on KD had considerably lower triglyceride levels than their sham KD counterparts, implying that the absence of ovarian hormones may improve lipid metabolism in females.

**Figure 4:**
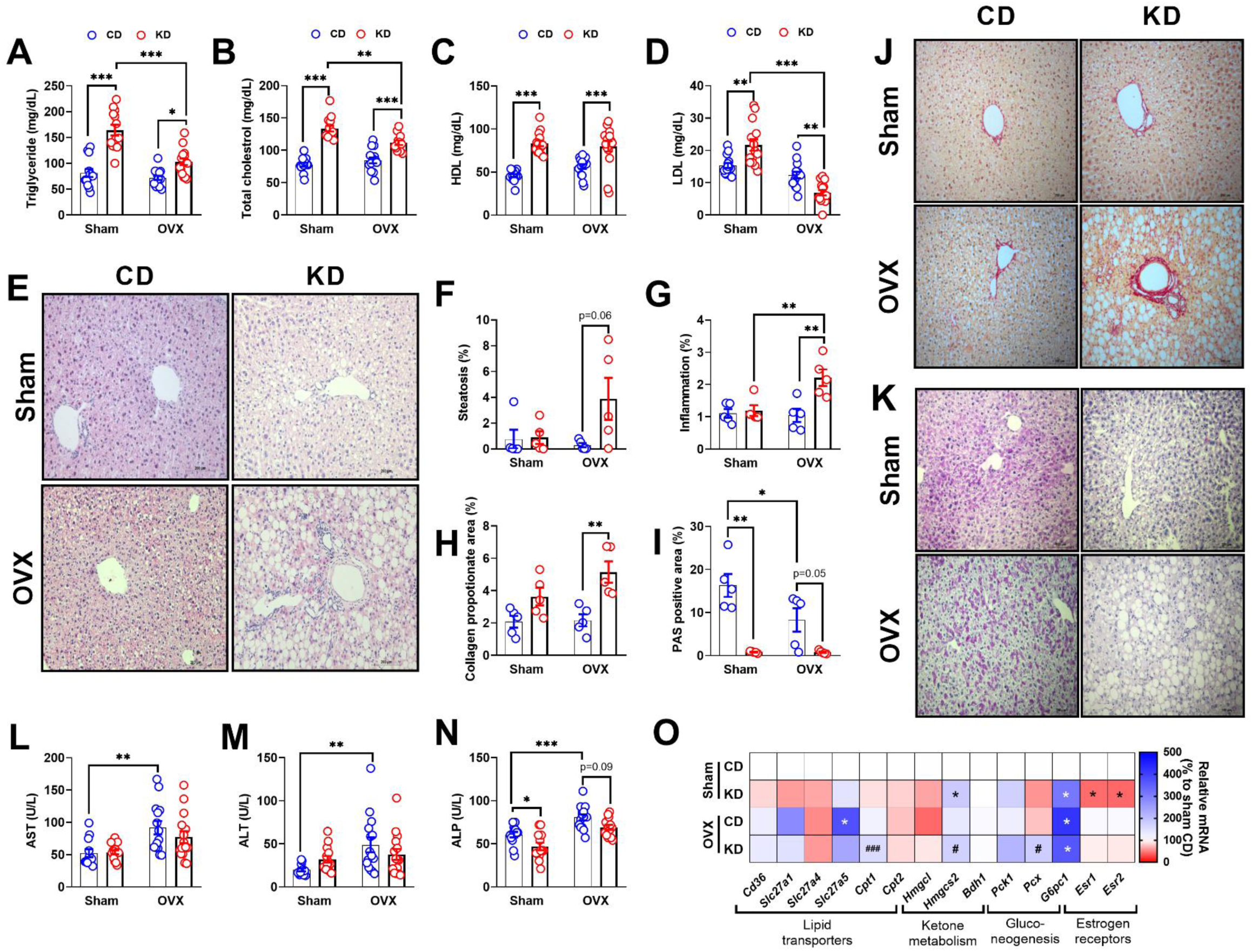
Effects of KD on plasma lipid levels, lipid metabolism and hepatic health in sham and OVX female mice. (**A–D**) Plasma lipid profile after 4 weeks of diet: **(A)** Triglycerides (mg/dL), **(B)** Total cholesterol (mg/dL), **(C)** HDL (mg/dL), and **(D)** LDL (mg/dL) levels. **(E)** Representative H&E-stained liver sections (Scale bar = 200 μm). **(F)** Steatosis score, **(G)** Inflammation score, **(H)** Collagen proportional area (%), and **(I)** PAS-positive area (%) quantification. **(J)** Representative Masson’s trichrome-stained liver sections showing collagen deposition (Scale bar = 200 μm). **(K)** Representative PAS-stained liver sections showing glycogen content (Scale bar = 200 μm). **(L–N)** Liver dysfunction marker enzymes: **(L)** AST (U/L), **(M)** ALT (U/L), and **(N)** ALP (U/L). **(O)** Relative mRNA expression of hepatic genes involved in lipid transport, ketone metabolism, gluconeogenesis, and estrogen signaling, normalized to Sham CD. Data are presented as mean ± SEM. Statistical significance: *p<0.05, **p<0.01, ***p<0.001, ****p<0.0001. Sample sizes: Panel A, n = 13 for Sham CD and Sham KD; n = 15 for OVX CD and OVX KD. Panel B, n = 13 for Sham CD; n = 14 for Sham KD; n = 15 for OVX CD; n = 13 for OVX KD. Panel C-D, n = 14 for Sham CD and Sham KD; n = 15 for OVX CD and OVX KD. Panel F-I, n = 5 per group. Panel L, n = 12 for Sham CD; n = 14 for Sham KD and OVX CD; n = 15 for OVX KD. Panel M, n = 12 for Sham CD; n = 14 for Sham KD; n = 15 for OVX CD and OVX KD. Panel N, n = 13 for Sham CD and Sham KD; n = 15 for OVX KD; n = 12 for OVC KD. Panel O, n = 6 per group. **CD**: Control diet; **KD**: Ketogenic diet; **OVX**: Ovariectomized; **Sham CD**: Group subjected to the sham surgery with a control diet; **Sham KD**: Group subjected to the sham surgery with a ketogenic diet; **OVX CD**: Group subjected to the OVX surgery with a control diet; **OVX KD**: Group subjected to the OVX surgery with a ketogenic diet; **H&E**: Hematoxylin and eosin; **PAS**: Periodic acid–Schiff; **HDL**: High-density lipoprotein; **LDL**: Low-density lipoprotein; **AST**: Aspartate aminotransferase; **ALT**: alanine aminotransferase; **ALP**: alkaline phosphatase.

Similarly, KD also induced hypercholesterolemia in sham and OVX females. Crucially, OVX females on KD had considerably lower cholesterol levels than their sham KD counterparts **(Fig. 4B)**. Plasma HDL-C levels were similarly higher in sham and OVX females on KD compared to their CD counterparts **(Fig. 4C)**. In contrast, LDL-C levels were significantly higher in sham females on KD compared to those given CD. However, in OVX females, KD reduced LDL levels when compared to CD counterparts **(Fig. 4D)**. Taken together, these findings indicate that while KD induces hyperlipidemia in both sham and OVX females, the lower LDL levels observed in OVX females on KD suggest that ovarian hormone loss alters lipid handling pathways, potentially leading to a more favorable redistribution of circulating lipids under high-fat dietary conditions.

### Metabolic adaptations to KD in females without molecular evidence of liver dysfunction

Building on our previous observation of KD-induced hyperlipidemia in sham and OVX females, we next examined hepatic consequences of KD and found steatosis, fibrosis, and metabolic adaptations at the histopathological levels without concurrent changes in molecular or pathological markers of liver dysfunction, both in sham and OVX female mice.

H&E staining revealed steatosis in the liver of OVX females on KD compared to those on CD, although the difference was not statistically significant **(Fig. 4E, F)**. However, KD suppressed the hepatic mRNA expression of peroxisome proliferator-activated receptor gamma (*Pparγ*) in sham and OVX females **(Fig. S2)**, suggesting that KD modulates hepatic lipid metabolism in females.

KD significantly enhanced the frequency of infiltrating immune cells around hepatic triads of OVX females **(Fig. 4E, G)**, implying that gonadal hormones play a key role in mediating KD-induced inflammation in the liver. In contrast, OVX increased mRNA expression of Interleukin 1 beta (*Il1β*) in the liver of females, while no such increase was observed after KD supplementation **(Fig. S2)**. Meanwhile, KD suppressed Interleukin 6 (*Il6*) transcription in sham females alone. The mRNA expression pattern of hepatic pro-inflammatory genes reveals that KD induces immune cell accumulation in the liver of OVX females, while inhibiting their pro-inflammatory response.

Furthermore, PSR staining revealed that KD promoted collagen deposition around the hepatic portal triads of OVX females **(Fig. 4H, J)**. However, despite the histological abnormalities, the mRNA expression of fibrosis-related genes, including tissue inhibitor of metalloproteinase 1 (*Timp1*), collagen type I alpha 1 chain (*Col1a1*), and transforming growth factor beta 1 (*Tgfβ1*), remained unchanged in those females **(Fig. S2)**. This suggests that, while the KD causes localized collagen accumulation in the liver of females, these changes are not accompanied by transcriptional activation of classical pro-fibrotic markers, implying that collagen deposition may be an early hepatic remodeling event by KD rather than an established fibrosis.

Hepatic PAS staining further revealed that KD reduced hepatic glycogen stores in both sham and OVX females. Furthermore, OVX alone was able to decrease the hepatic glycogen levels in females regardless of diet **(Fig. 4I, K)**, suggesting that both the absence of ovarian hormones and the KD can decrease hepatic glycogen stores in females.

A considerable increase in liver dysfunction marker enzymes such as AST **(Fig. 4L)**, ALT **(Fig. 4M)** and ALP **(Fig. 4N)** was observed in the plasma of OVX females on CD. However, no such increase in AST, ALT and ALP levels was observed in OVX females supplemented with KD, suggesting that while ovarian hormone deprivation exacerbates liver damage in females, KD somehow prevents this damage.

### KD differentially modulates metabolic and estrogen receptor genes in the liver of sham vs. OVX females

To further understand the impact of KD on hepatic ketone and glucose metabolism, we examined the mRNA expression of rate-limiting genes for ketogenesis, ketolysis and gluconeogenesis in hepatic tissue of sham and OVX females **(Fig. 4O)**.

The mRNA expression of hepatic fatty acid transport protein 5 (FATP5, *Slc27a5)* was elevated in OVX females on CD. However, KD prevented the elevation of *Slc27a5* in OVX females. No change in the transcription of other fatty acid transporters, including *Slc27a1 (*FATP1), *Slc27a4* (FATP4), and *Cd36,* was observed in any of the experimental groups. Furthermore, we assessed transcript levels of genes implicated in the carnitine shuttle system, such as Carnitine Palmitoyltransferase 1 (*Cpt1*) and *Cpt2*, which play an important role in fatty acid oxidation by enabling fatty acid transport from cytosol to the mitochondrial matrix. KD increased transcript levels of hepatic *Cpt1* only in OVX females. In contrast, there were no discernible differences in *Cpt2* mRNA expression among the experimental groups. The mRNA expression of 3-hydroxy-3-methylglutaryl-CoA synthase 2 (*Hmgcs2*), the rate-limiting enzyme for ketogenesis, was considerably higher in sham and OVX females on KD than in their CD counterparts. In contrast, no alterations were seen in the transcription of the 3-hydroxy-3-methylglutaryl-CoA lyase (*Hmgcl*) and 3-hydroxybutyrate dehydrogenase 1 (*Bdh1*) genes across any of the treatment groups. These findings indicate that KD influences the transcription of the fatty acid transporters (*Slc27a5, Cpt1*) and the rate-limiting enzyme for ketogenesis (*Hmgcs2*) in females, depending on their gonadal hormone status.

KD also increased transcription of hepatic glucose-6-phosphatase (*G6pc1*), the rate-limiting enzyme located in the endoplasmic reticulum, which catalyzes the final step of gluconeogenesis by producing glucose from glucose-6-phosphate, in both sham and OVX females. However, KD increased transcription of hepatic pyruvate carboxylase (*Pcx*), a crucial mitochondrial enzyme that catalyzes the first step of gluconeogenesis by converting pyruvate to oxaloacetate, only in OVX females. Overall, these findings indicate that KD may promote hepatic gluconeogenesis in females regardless of gonadal hormonal state and that pyruvate may be the key precursor for gluconeogenesis in OVX females.

KD also decreased transcription of hepatic *Esr1* and *Esr2* estrogen receptors in sham but not in OVX females, implying that KD impairs hepatic estrogen signaling in ovaries-intact females.

### KD increases biochemical markers of renal dysfunction and induces electrolyte imbalance in female mice

To evaluate the impact of a KD on renal function and electrolyte balance, we assessed markers of nitrogen and electrolyte balance in the plasma of sham and OVX female mice **(Fig. S3)**. KD decreased plasma urea levels in sham females while increasing blood urea in OVX females compared to their CD counterparts **(Fig. S3A).** These findings suggest that KD exerts differential effects on renal physiology and nitrogen metabolism in females depending on their gonadal hormone status.

Plasma electrolyte and protein analysis revealed that KD increased potassium levels in sham female mice, not in OVX females **(Fig. S3B)**. However, plasma sodium and chloride concentrations were significantly increased in OVX females regardless of diet **(Fig. S3 C-D)**. Likewise, albumin levels were also elevated in OVX females on CD, while KD prevented the hyperalbuminemia in those females **(Fig. S3F)**. The simultaneous rise in electrolytes and protein fractions suggests a possible concentration effect, most likely reflective of dehydration or altered fluid balance in OVX females.

Together, these findings indicate that KD not only perturbs nitrogen metabolism but also induces electrolyte imbalance in females, depending on their gonadal hormone status, highlighting a higher susceptibility of gonadal hormone-deficient females to renal and systemic fluid imbalance under ketogenic conditions.

### KD induces low-grade inflammation in sham, while preserving immune functions in OVX female mice

To investigate the immunomodulatory effects of KD, we performed a cytokine and chemokine array analysis in the plasma of sham and OVX female mice **(Fig. 5)**. Sham females maintained on KD displayed a broad elevation of pro-inflammatory mediators compared to their CD counterparts. Notably, circulating levels of cytokines IL-5, IL-6, IL-7, IL-17A, IL-28, IL-33, and IFN-γ were increased, alongside an increase in chemokines such as CCL5 and CXCL2. In addition, KD-fed sham females showed altered profiles of secretory proteins and growth factors linked to inflammation and vascular damage, including a rise in PD-ECGF, VEGF, CD-40, osteoprotegerin, CD14, fetuin, endostatin, and soluble LDL R. These results collectively point to a state of low-grade inflammation following KD in sham females.

**Figure 5.**
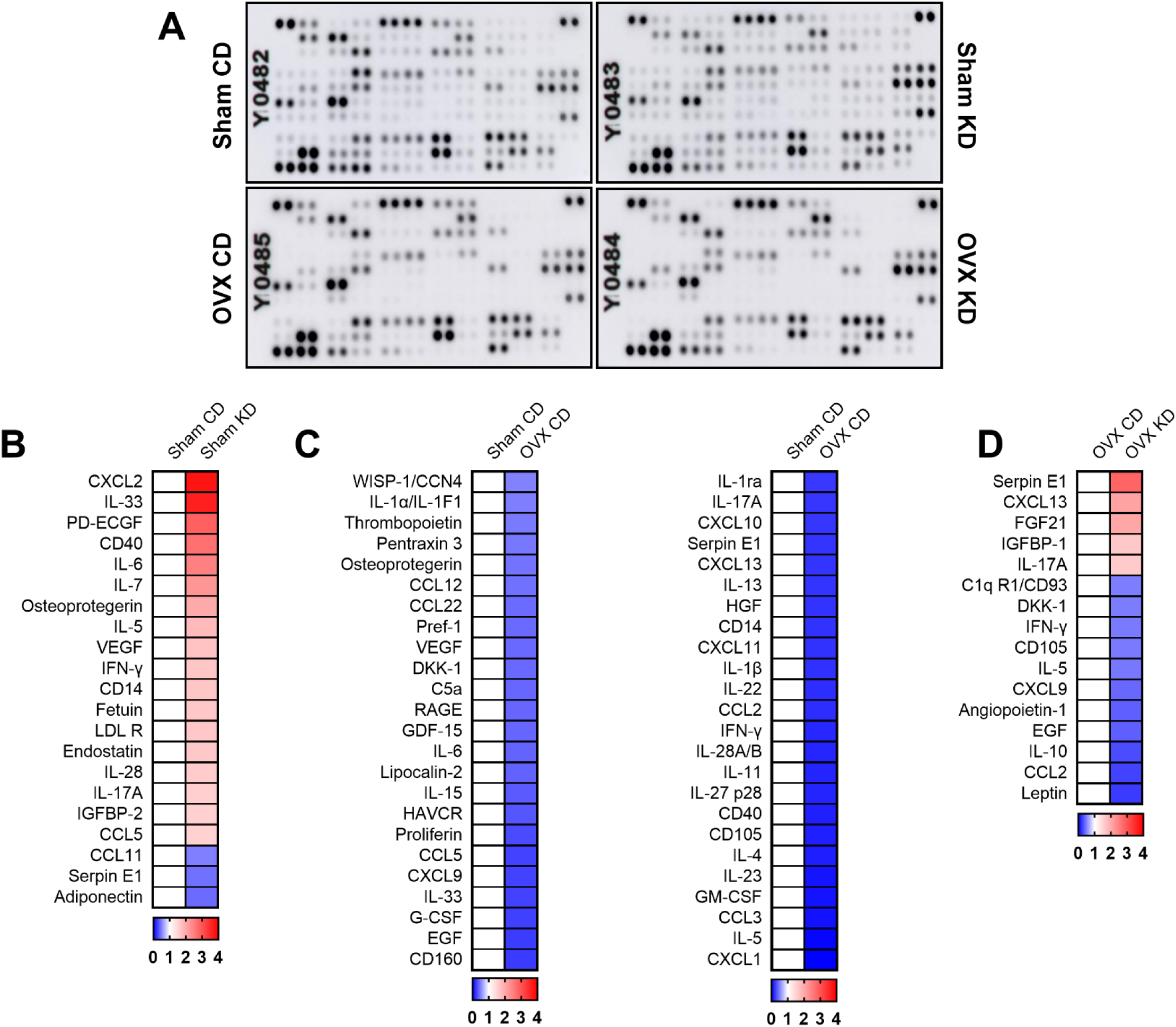
Effects of KD on systemic cytokine, chemokine and growth factor profiles of sham and OVX female mice. (**A–D**) Multiplex cytokine array analysis of plasma from sham and OVX female mice maintained on either CD and KD for 4 weeks. **(A)** Representative images of dot blots developed from each group. Heatmaps showing relative fold changes in indicated analytes between (**A**) Sham CD vs. Sham KD, (**B**) Sham CD vs. OVX CD, and (**D**) OVX CD vs. OVX KD. Each row represents an individual protein; red denotes higher and blue denotes lower expression (scale 0–4). Sample size: Panel A-D, n =14 samples pooled to generate a single composite sample per group. **CD**: Control diet; **KD**: Ketogenic diet; **OVX**: Ovariectomized; **Sham CD**: Group subjected to the sham surgery with a control diet; **Sham KD**: Group subjected to the sham surgery with a ketogenic diet; **OVX CD**: Group subjected to the OVX surgery with a control diet; **OVX KD**: Group subjected to the OVX surgery with a ketogenic diet.

Compared to sham controls, OVX females displayed a marked reduction in basal pro-inflammatory cytokines, chemokines and other inflammatory factors. This broad downregulation of immune mediators indicates a suppressed systemic immune tone in OVX females, suggesting that ovarian hormone deficiency blunts basal immune activity. In contrast, KD preserved systemic immune functions in OVX females, as indicated by elevated levels of serpin E1, CXCL13, FGF21, IGFBP-1 and IL-17A. These observations highlight a striking divergence in the immunomodulatory effects of KD between sham and OVX females. While KD promotes a state of low-grade inflammation in sham females, it paradoxically preserves or partially restores immune signaling in OVX females. This sex-hormone–dependent dichotomy underscores the critical role of ovarian hormones in shaping systemic immune responses to dietary interventions such as KD.

### KD differentially modulate the gut microbiota of sham and OVX female mice

To determine whether the physiological, metabolic and neurological effects of KD are mediated through changes in the gut microbiota, we performed 16S rRNA gene sequencing on DNA isolated from fecal samples of sham and OVX female mice maintained on KD or CD. Alpha diversity, reflecting within-sample microbial richness, was assessed using the observed features **(Fig. 6A)** and Shannon index **(Fig. 6B)**. KD significantly increased microbial diversity in sham females (p=0.008) compared to their CD counterparts, indicating an expansion of gut microbial richness under KD. In contrast, OVX females on KD exhibited reduced observed-feature alpha diversity relative to OVX females on CD (p=0.008) and their sham KD counterparts (p=0.008).

**Figure 6:**
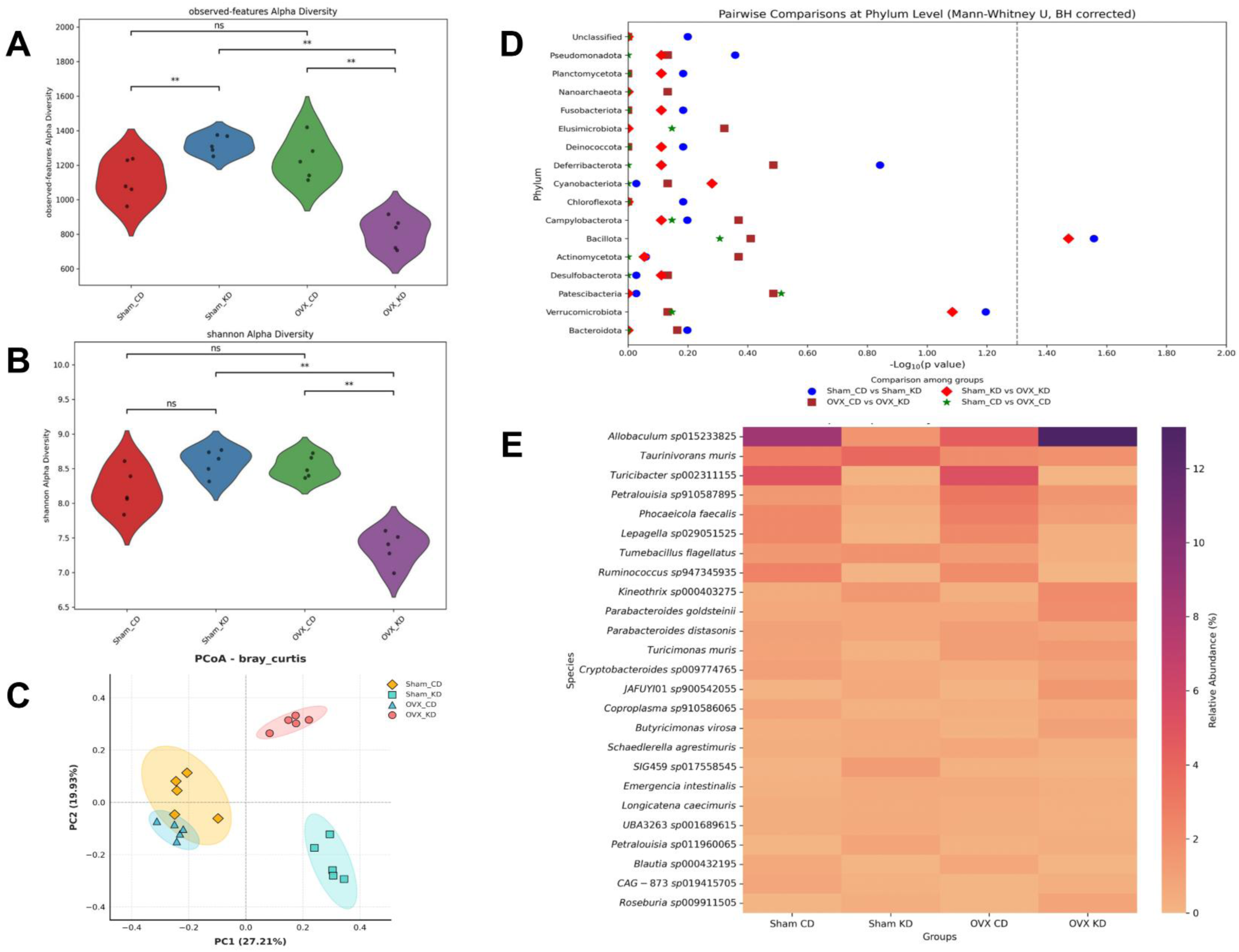
Effect of KD on gut microbiota of sham and OVX female mice. (A-B) Violin plots showing alpha diversity of the gut microbiome across the four experimental groups. **(A)** Observed features (species richness) and **(B)** Shannon diversity index (richness and evenness). Significance between groups is indicated as: ns: not significant; **: p < 0.01. **(C)** Principal Coordinate Analysis (PCoA) of Bray-Curtis Dissimilarity demonstrating Beta diversity among the four experimental groups, with PC1 (27.21%) and PC2 (19.93%). Samples cluster distinctly by group, and PERMANOVA confirmed significant differences in microbial diversity (pseudo-F = 5.386, p = 0.001). **(D)** Pairwise Comparisons at the phylum level. Dot-plot representing pairwise comparisons of microbial relative abundance at the phylum level among experimental groups using the Mann-Whitney U test with Benjamini-Hochberg adjustment. The x-axis represents the -log10(p-value) of relative abundance, and the y-axis represents the phyla present. Colors and symbols indicate the specific pairwise comparison: Blue circle: Sham CD vs Sham KD; Red Diamond: Sham KD vs OVX KD; Brown square: OVX CD vs OVX KD; Green star: Sham CD vs OVX CD. **(E)** Heatmap of top 25 species based on relative abundance. Heatmap showing the relative abundance (%) of the top 25 bacterial species across four experimental groups: Sham CD - control diet; Sham KD - ketogenic diet; OVX CD - ovariectomized, control diet; and OVX KD - ovariectomized, ketogenic diet. The color intensity represents the relative abundance of each species, with darker shades indicating higher abundance.

Shannon index analysis revealed no significant differences in microbial diversity between sham CD and sham KD groups (p = 0.056) or between sham CD and OVX CD groups (p = 0.095). However, OVX females maintained on KD exhibited a significant reduction in Shannon diversity compared to their CD and sham KD counterparts (p = 0.008), suggesting that ovarian hormone loss alters the microbial response to KD and may limit its diversity-promoting effects.

To further evaluate group-level differences in microbial community composition, beta diversity was assessed using Bray–Curtis dissimilarity and visualized through Principal Coordinate Analysis (PCoA) **(Fig. 6C)**. The PCoA plot demonstrated clear separation of samples across experimental groups, which was statistically supported by PERMANOVA (p = 0.001), confirming a significant influence of KD on gut microbial composition of sham and OVX females. The first two principal coordinates (PC1 and PC2) together accounted for 47.14% of the total variance.

Pairwise comparison of phylum-level relative abundances, performed using the Mann–Whitney U test with Benjamini–Hochberg correction, demonstrated a significant alteration in gut microbial composition in response to KD in both sham and OVX females **(Fig. 6D)**. Specifically, KD feeding led to a marked increase in the relative abundance of the Bacillota phylum in sham females compared to those maintained on a CD. A similar elevation in Bacillota abundance was observed in KD-fed OVX females relative to their sham KD counterparts. These findings suggest that KD induces an enrichment of Bacillota in the gut microbiota of females, an effect that appears to occur independently of gonadal hormone status.

A heatmap of the top 25 microbial species across all samples revealed clear clustering patterns and marked differences in relative abundance among the experimental groups **(Fig. 6E)**. Dominant species included Allobaculum sp015233825, Taurinivorans muris, and Turicibacter sp002311155, although their distribution varied substantially between sham and OVX females on KD. The KD prompted a distinct species-level reorganization of the gut microbiota, characterized by increases in certain taxa and corresponding decreases in others relative to CD-fed counterparts in both sham and OVX mice. These findings underscore a strong interactive effect of diet and hormonal status on gut microbial composition, suggesting that although a core set of species remains dominant across groups, their relative abundances are dynamically shaped by KD and ovarian hormone deficiency.

Pearson correlation analysis was performed to evaluate the associations between microbial species, physiological measures, and behavioral outcomes of KD in sham and OVX female mice **(Fig. 7)**. In sham females, several microbial species, including *Petralouisia sp011960065*, *Alistipes sp021204515*, *JAFUYI01 sp900542055*, *Emergencia intestinalis*, *Kineothrix sp000403275*, *Butyricimonas virosa*, *Taurinivorans muris*, *Petralouisia sp910587895*, *Coproplasma sp910586065*, *Turicimonas muris*, and *Phocaeicola faecalis*, showed significant correlations with metabolic and behavioral parameters. In contrast, in OVX females, correlations were observed between *Cryptobacteroides sp009774765*, *Coproplasma sp910586065*, *Parabacteroides goldsteinii*, *Schaedlerella agrestimuris*, and *Tumebacillus flagellatus* with metabolic and behavioral outcomes. Overall, the correlation patterns suggest that distinct microbial species may modulate the physiological and behavioral effects of KD depending on the ovarian hormonal status of the females. These findings highlight the influence of ovarian hormones in shaping the gut microbiota–metabolism–brain axis under ketogenic dietary conditions.

**Figure 7:**
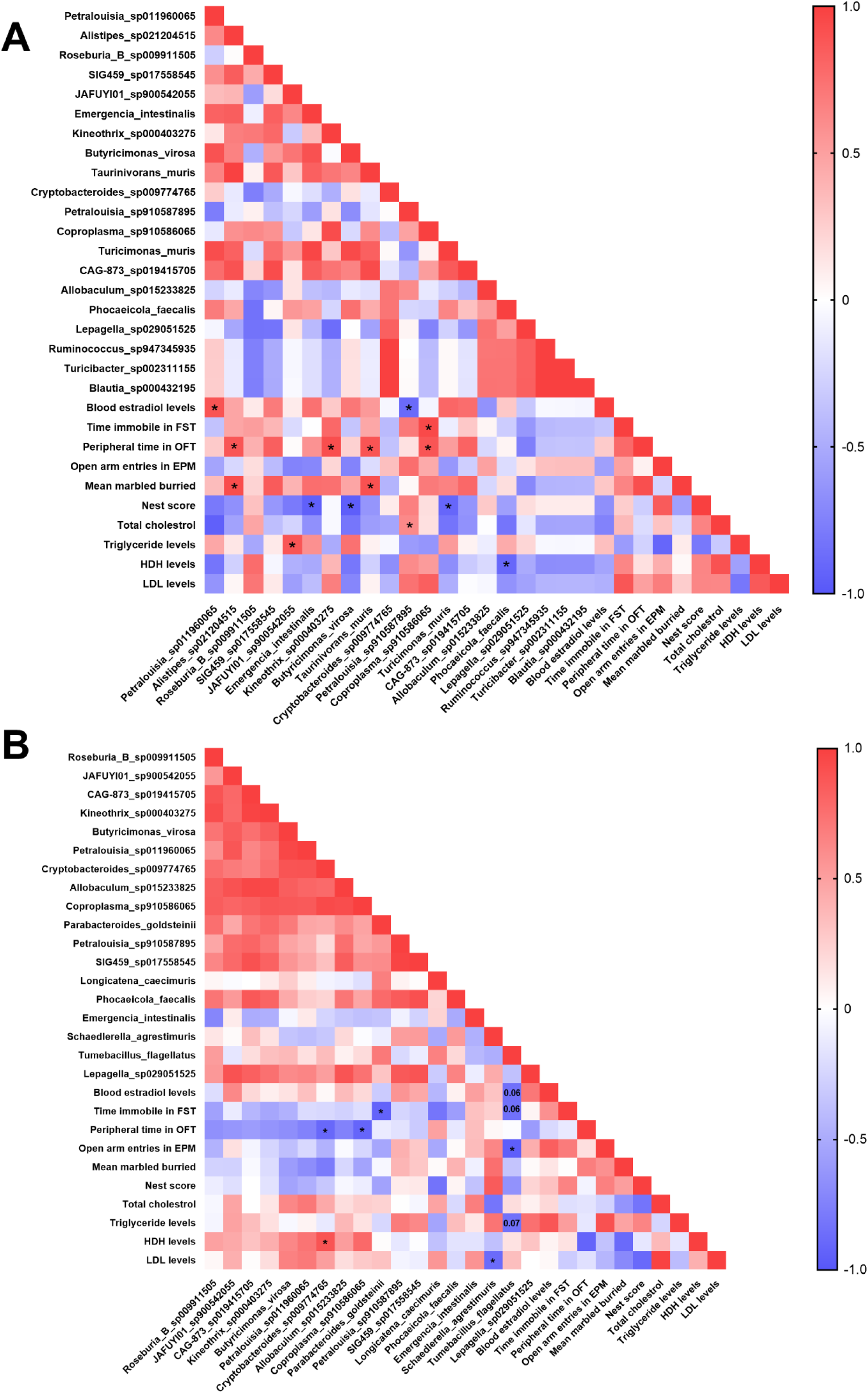
Pearson Correlation Matrix of gut microbial species abundance, physiological, metabolic and behavioral outcomes in KD-fed sham **(A)** and OVX **(B)** female mice. The correlation matrix is based on Pearson’s linear correlation. Each square is a correlation with n=5 mice/group.

**Figure S1.**
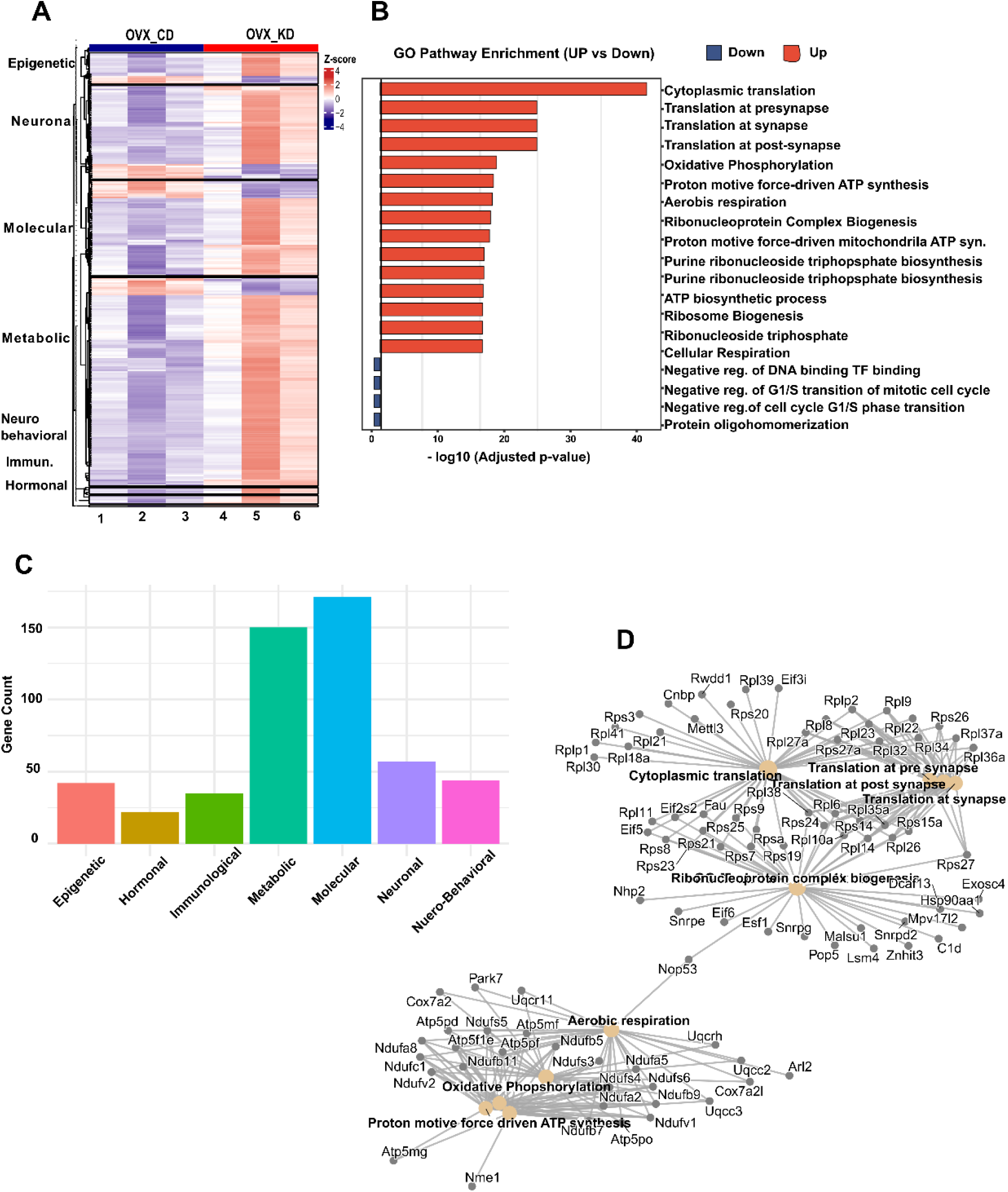
Integrated visualization of GO-module enrichment and connectivity. **(A)** Module-specific heatmap showing Z-score–scaled DEGs across seven functional modules, with consistent upregulation of genes in OVX KD and downregulation in OVX CD groups. **(B)** GO enrichment bar plot ranks top up- and downregulated pathways, highlighting strong activation of translation, synaptic, and oxidative phosphorylation processes. **(C)** Bar plot of module-wise gene counts showing dominance of metabolic and molecular categories in response to KD, supporting multi-domain transcriptional activation under ovarian hormone deprivation. **(D)** cnetplot network links DEGs to enriched GO terms, revealing shared genes bridging translational and mitochondrial pathways.

**Figure S2:**
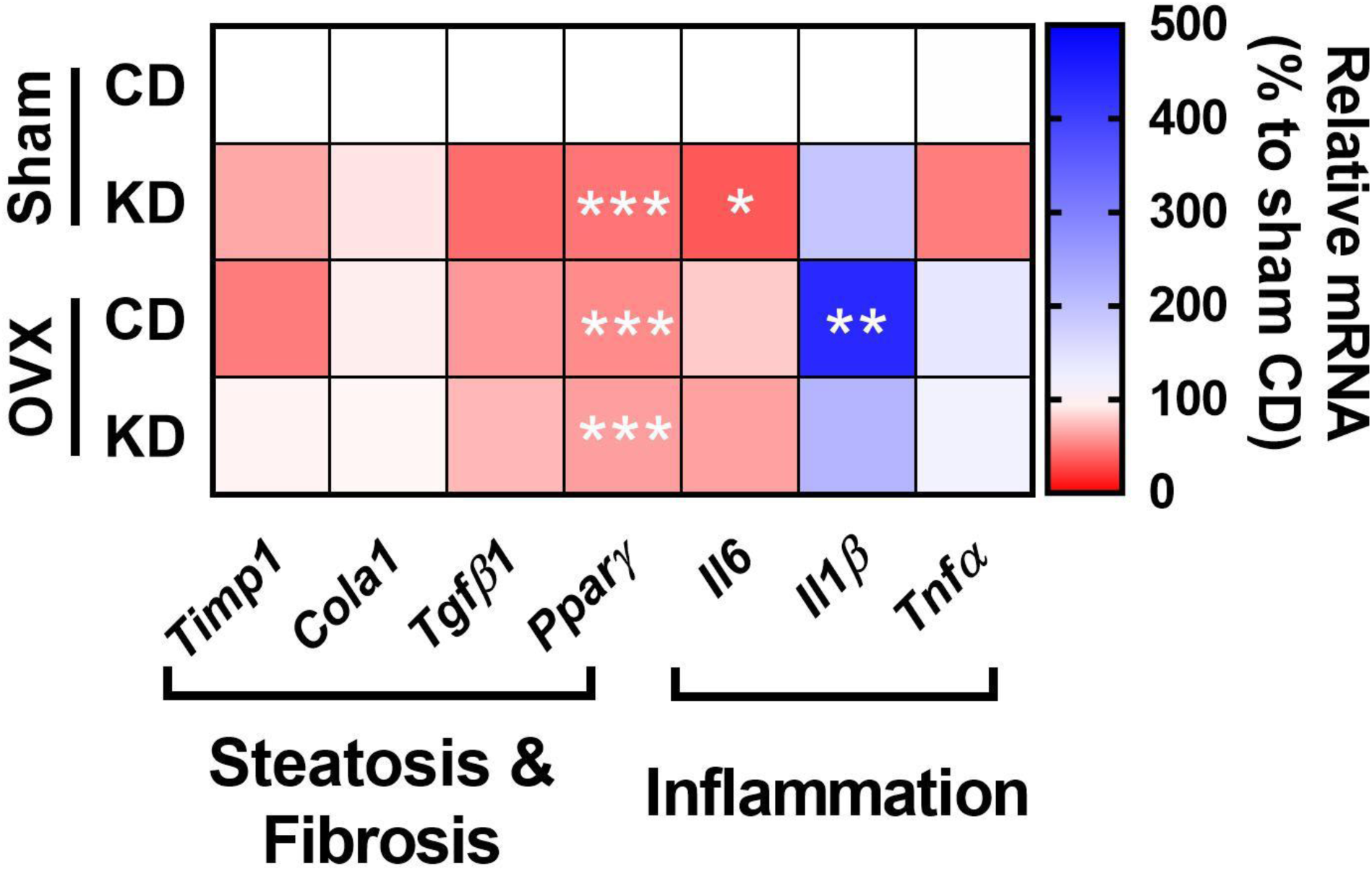
Transcriptional profiling of liver fibrosis, steatosis, and inflammation marker genes in response to KD. Relative mRNA expression of hepatic genes associated with steatosis, fibrosis, and inflammation, normalized to sham CD. All data are expressed as mean ± SEM. Statistical significance is indicated as *p<0.05, ** p<0.01, *** p<0.001, and **** p<0.0001. Sample size: n = 6 per group. **CD**: Control diet; **KD**: Ketogenic diet; **OVX**: Ovariectomy; **Sham CD**: Group subjected to the sham surgery with a control diet; **Sham KD**: Group subjected to the sham surgery with a ketogenic diet; **OVX CD**: Group subjected to the OVX surgery with a normal control diet; and **OVX KD**: Group subjected to the OVX surgery with a ketogenic diet.

**Figure S3:**
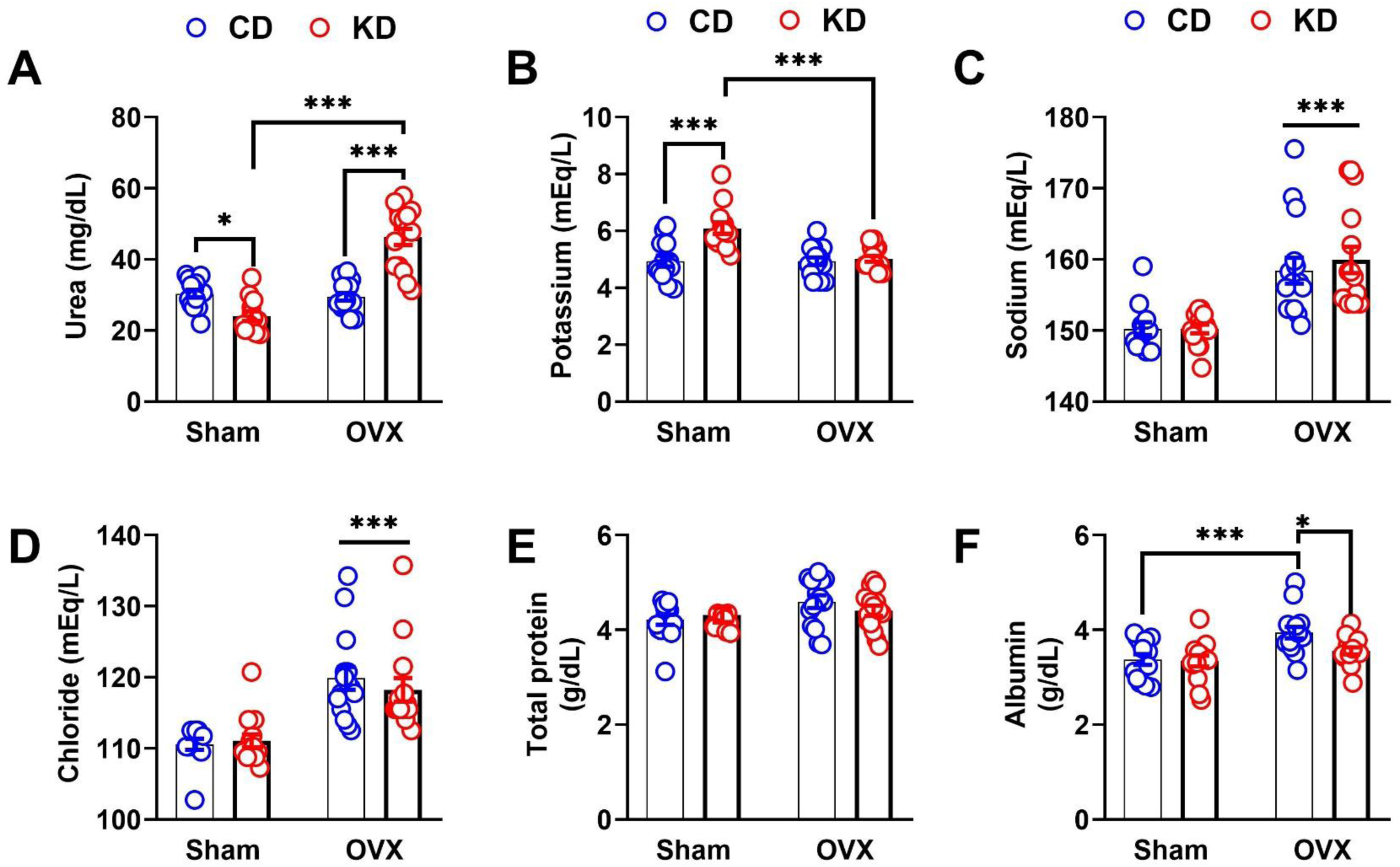
Effect of KD on kidney dysfunction markers and electrolyte levels in shama nd OVX female mice. **(A)** Urea (mg/dL). (**B-D**) Plasma electrolyte level: (**B**) Potassium (mEq/L), (**C**) Sodium (mEq/L), (**D**) Chloride (mEq/L). (**E**) Total protein (g/dL). (**F**) Albumin (g/dL). All data are expressed as mean ± SEM. Statistical significance is indicated as *p<0.05, ** p<0.01, *** p<0.001, and **** p<0.0001. Sample size Panel A, n = 14 for Sham CD; n = 13 for Sham KD; n = 15 for OVX CD and OVX KD Panel B-C, n = 13 for Sham CD; n = 14 for Sham KD; n = 15 for OVX CD and OVX KD Panel D, n = 12 for Sham CD; n = 15 for Sham KD; n = 15 for OVX CD and OVX KD Panel E-F, n = 14 for Sham CD and Sham KD; n = 15 for OVX CD and OVX KD **CD**: Control diet; **KD**: Ketogenic diet; **OVX**: Ovariectomy; **Sham CD**: Group subjected to the sham surgery with a control diet; **Sham KD**: Group subjected to the sham surgery with a ketogenic diet; **OVX CD**: Group subjected to the OVX surgery with a control diet; and **OVX KD**: Group subjected to the OVX surgery with a ketogenic diet.

## Discussion

The current study demonstrates that KD differentially affects metabolic and hormonal parameters in female mice, depending on their ovarian hormone status. For example, OVX mice on CD had higher body weight, even with reduced caloric intake, in agreement with previous studies that highlight the role of estrogen in maintaining metabolic homeostasis and body weight in female mice [19]. However, in contrast to the earlier study in male mice [20], KD did not decrease food intake in female mice, regardless of ovarian hormone status, suggesting that KD may promote weight loss in females by boosting their energy expenditure. Notably, KD decreased the circulating estradiol levels in sham females, aligning with recent evidence that prolonged KD may suppress estrogen levels in young female mice[21] and middle-aged obese women[22]. Such estrogen modulation by KD highlights an important caveat in applying ketogenic interventions in young females, particularly regarding reproductive hormone balance. Despite significant metabolic shifts, serum corticosterone levels remained unchanged, arguing against hypothalamic-pituitary-adrenal (HPA) axis involvement and supporting the specificity of dietary and hormonal interventions in driving observed outcomes.

KD feeding confers broad behavioral resilience in OVX female mice, spanning anxiety-like, depressive-like, and compulsive domains. Across multiple validated paradigms, including the open field, elevated plus maze, forced swim, marble burying, and nest building tests, KD consistently shifted behavior toward reduced anxiety, compulsivity and depressive-like phenotypes while promoting overall well-being. These robust effects indicate that estrogen depletion creates a neurobiological context in which KD’s actions on brain energy metabolism and synaptic function translate into measurable improvements in affective behavior. Transcriptomic analyses of the mPFC provide convergent molecular support for these behavioral gains. KD induced a clear metabolic reprogramming of the mPFC characterized by marked downregulation of *Pfkfb3* transcripts, a key glycolytic rate-controlling enzyme, and strong upregulation of *Txnip*, a sensor of cellular energy status and a negative regulator of cellular glucose uptake [23,24]. Together, these transcriptomic shifts point to reduced glucose metabolism in the mPFC and increased reliance on fatty acid or ketone oxidation. This metabolic redirection was further reinforced by coordinated upregulation of multiple electron transport chain (ETC) and ATP synthase subunits associated genes, suggesting an enhanced mitochondrial capacity to support sustained oxidative phosphorylation under ketone-based fueling. Such enhancements in mitochondrial efficiency and substrate flexibility are well aligned with the neuroprotective and stress-buffering properties attributed to KD [25–27]. At the synaptic level, KD-driven increases in *Cabp1*, *Syt11*, and *Stmn3* gene expression suggest strengthened calcium buffering, improved vesicle trafficking, and enhanced structural plasticity. These changes are consistent with the synapse-stabilising and plasticity-enhancing effects previously attributed to KD [35]. They may represent a key mechanistic link between KD-driven metabolic reprogramming and the behavioral resilience observed in OVX females. Together, the transcriptional profile points to an mPFC network that is more energetically efficient, structurally supported, and capable of maintaining stable activity patterns under conditions that typically elicit vulnerability following estrogen loss. In summary, our results provide compelling evidence that KD exerts powerful behavioral and molecular benefits in OVX female mice by reconfiguring mPFC metabolic networks and reinforcing synaptic resilience.

KD increased circulating triglyceride and total cholesterol levels in both sham and OVX mice. Notably, the extent of hyperlipidemia was less pronounced in the OVX females compared to their sham counterparts, suggesting that gonadal hormone pre-deficiency prevented KD-induced hyperlipidemia in females. In the present study, the marked reduction in systemic estradiol levels and hepatic expression of its receptors (*Esr1* and *Esr2*) likely impairs lipid transport and the metabolic capacity of hepatocytes, contributing to KD-induced hyperlipidemia in sham females, as the availability of estrogen and its cognate receptors is inversely proportional to dyslipidemia [28,29]. In line with this, KD also increased LDL levels in sham females, pointing toward enhanced atherogenic potential. This elevation coincided with a reduction in circulating adiponectin, a hormone known to modulate LDL metabolism [30], implying that adiponectin suppression may have contributed to the LDL cholesterol imbalance. Conversely, the sustained expression of hepatic estrogen receptors and circulating adiponectin in OVX females may have protected against KD-induced hyperlipidemia by preserving hepatic lipid transport and metabolism, though further work is needed to clarify how KD influences hepatic estrogen signaling and its downstream metabolic processes.

Recent studies have reported that prolonged KD can induce hepatic inflammation and steatosis by disrupting insulin signaling [31,32]. However, some studies have described the hepatoprotective effects of KD by attenuating oxidative stress, mitochondrial biogenesis, and fibrosis [33,34]. Consistent with this duality, OVX mice on KD exhibited histological signs of inflammation, steatosis, and early fibrosis without significant transcriptional upregulation of pro-inflammatory or fibrotic marker genes. This pattern likely reflects an initial adaptive response of the liver to KD rather than hepatic pathological progression. Further PAS staining and elevated *Hmgcs2* mRNA expression also supported the metabolic adaptation to KD, as evidenced by reduced hepatic glycogen stores and elevated circulating ketone bodies [35,36]. KD also promoted hepatic gluconeogenesis, as evidenced by increased *G6pc1* mRNA expression, independently of ovarian hormonal status. This finding is consistent with earlier work demonstrating induction of gluconeogenesis by KD, which further helps to normalise blood glucose levels despite carbohydrate restriction [37,35]. Elevated levels of liver dysfunction enzymatic markers, such as ALT, AST, and ALP, were observed in OVX females on CD, consistent with previous findings highlighting the crucial role of estrogen in maintaining liver health [38]. However, these markers were not elevated in OVX females on KD, suggesting that KD may prevent OVX-induced liver dysfunction in females. This aligns with preclinical evidence showing that KD improves liver functions by enhancing mitochondrial dynamics and β-oxidation [39]. Overall, these findings indicate that KD induces nutritional ketosis, largely independent of ovarian hormonal milieu. However, KD may elevate the risk of atherogenic processes in young females, underscoring the need for cautious interpretation and longitudinal assessment in translational contexts.

Estrogen deprivation markedly elevates the risk of chronic kidney disease, as evidenced by increased blood urea and albumin levels, biomarkers of impaired glomerular filtration rate (GFR) in OVX rats [40]. Prior studies have further revealed that OVX-induced electrolyte disturbances originated from altered renal signaling, specifically the dysregulation of endothelin-1, purinergic (P2) receptors, and G-protein coupled estrogen receptors [41,42], collectively contributing to a higher risk of hypertension in post-menopausal females. While KD has been shown to have nephroprotective effects in genetic and acquired models of kidney disease, such as cystinosis and polycystic kidney disease [43,44]. The present data indicate that KD after OVX increased serum urea levels in this biologically vulnerable state. These findings suggest that KD post-menopause may further elevate the risk of metabolic acidosis, impairing glomerular filtration rather than ameliorating kidney dysfunction in gonadal hormone-deficient females. Consistent with clinical guidelines, confirmation of renal impairment should include assessment of plasma creatinine alongside urea, as simultaneous elevations are strong indicators of chronic kidney disease or acute kidney injury. Therefore, future studies should incorporate creatinine measurements or eGFR to evaluate KD-induced renal outcomes in OVX models.

KD is increasingly recognized for its capacity to modulate innate and adaptive immune responses across preclinical and clinical settings [5,4,45] by regulating NLRP3 and FGF21 signaling [46] and reshaping gut microbiota composition [47]. Interestingly, KD has been reported to act as a primary stimulus for trained immunity, reprogramming innate immune cells to exhibit heightened responsiveness upon secondary challenge through a palmitic acid–ceramide axis [48]. These findings align with our observation of elevated pro-inflammatory markers in sham females on KD, suggesting that KD may establish a persistent immune imprint independent of metabolic disease states. However, when administered post-OVX, KD appeared to restore several immune and metabolic mediators that were otherwise suppressed in OVX females. Notably, KD upregulated protective factors such as FGF21 and IGFBP-1 and shifted the chemokine/cytokine milieu toward an anti-inflammatory state with reduced levels of IFN-γ, CXCL13, CCL2, CXCL9 and CD93. These adaptations are consistent with recent studies showing that KD dampens chronic inflammation and enhances metabolic resilience by regulating ketone metabolism and FGF21-mediated signaling [46,14]. Collectively, these findings suggest that KD can reprogram immune-metabolic interactions in sham and OVX females, conferring low-grade inflammation in young females while providing anti-inflammatory and metabolic advantages to gonadal hormone-deprived females. However, future studies are needed to determine whether KD during the reproductive stage can induce trained immunity and predispose to having a maladaptive inflammatory response to later-life challenges.

KD is recognized for its profound influence on gut microbiota composition, often leading to shifts in microbial diversity and the relative abundance of various phyla, which can subsequently impact host metabolism and overall health [49,47]. Studies have demonstrated that KD can significantly increase the proportion of firmicutes and elevate the firmicutes/bacteroidetes ratio [50]. Our findings corroborate these established patterns, as both sham and OVX mice on a KD exhibited altered diversity, with a notable enrichment of the phylum bacillota, formerly known as firmicutes. This phenomenon is likely attributable to the absence of carbohydrate content in KD, essential for many gut bacteria, leading to a decrease in overall microbial diversity [51]. This reduction in microbial load within the OVX KD group, despite maintaining diversity, might indicate a more profound impact on overall microbial biomass, potentially due to altered metabolic demands or hormonal influences in OVX subjects on KD. This observation suggests a complex interplay between diet, hormones, and microbial ecosystem, where diversity is maintained, but the absolute abundance of microorganisms is altered [52]. KD altered the relative abundance of the top 25 gut microbial species compared to CD across sham and OVX groups. The majority of species demonstrated compositional stability with only minor changes observed between diet and hormonal conditions; however, two species, *Allobaculum sp015233825* and *Taurinivorans muris,* displayed clear, diet-dependent shifts in both groups. These results are consistent with emerging evidence that KD exerts targeted effects on specific microbial populations [47,2]. *Allobaculum* abundance was increased in the OVX KD group, despite a drop in overall microbiota alpha diversity. This genus has been shown to modulate lipid metabolism by increasing the expression of angiopoietin-like 4 (ANGPTL4) in the gut of mice treated with a high-fat diet [53]. Further correlation analyses revealed that several gut microbial species were significantly associated with behavioral and metabolic parameters in KD-fed groups. Notably, the correlation patterns differed between OVX and sham females, indicating that gonadal hormones modulate how gut microbes influence host behavior and metabolism during KD. These findings underscore the multifaceted interplay among diet, microbial composition, and endocrine factors in regulating host physiology and behavior.

## Conclusion

In summary, this study identifies gonadal hormones as a critical regulator of KD adaptability in females. In gonadal hormone-sufficient females, KD exhibited estrogen loss, dyslipidemia, low-grade inflammation and suppressed hepatic estrogen signaling, whereas KD promoted behavioral and metabolic resilience while maintaining immune balance in gonadal hormone-deprived females. These findings demonstrate that estrogen signaling fundamentally shapes the metabolic, cognitive, and immunological outcomes of KD and underscore the importance of hormone-specific dietary strategies for women.

## Materials and methods

### Experimental animals and care

Adult female C57BL/6J mice (6–8 weeks old) were used for the study. Animals were procured from the Central Animal Facility (CAF) at the National Institute of Pharmaceutical Education and Research (NIPER), Mohali, India. Mice were housed in the Animal Experimentation Facility of the BRIC–National Agri-Food and Biomanufacturing Institute (NABI), Mohali, India, under specific pathogen-free conditions at 25 ± 2 °C with a 12 h light/dark cycle, and given free access to standard chow and water. Four to five mice were housed per individually ventilated cage (IVC) and acclimatised for 2 weeks before the experiment began. All procedures were approved by the Institutional Animal Ethics Committee (IAEC) of NABI and conducted in accordance with the guidelines of the Committee for the Control and Supervision of Experiments on Animals (CCSEA), Government of India.

### Surgery and dietary intervention

A total of 60 animals were recruited for the current study. Animals were randomly allocated into two groups: the ovariectomy (OVX) group (n = 30) and the sham-operated control group (n = 30). A bilateral OVX procedure was performed to deplete gonadal hormones using the method described previously [54], with slight modifications. Briefly, for the surgical induction of OVX, animals were initially anesthetized with 3% isoflurane and maintained under 1% isoflurane throughout the procedure. The abdominal area was shaved and disinfected with povidone-iodine solution. A small midline incision was made to open the abdominal cavity. The right uterine horn was identified, tied at its distal end using absorbable suture material, and the ovary was removed. The same procedure was repeated on the left side. The peritoneal wall was closed with absorbable sutures, and the skin was sealed using non-absorbable sutures. Sham surgeries were performed similarly but without the removal of the ovaries. After surgery, all animals received gentamycin (5 mg/kg; IP) and diclofenac (10 mg/kg, IP) to prevent infection and manage pain. Animals were housed individually for one week to allow postoperative recovery before being returned to their home cages. Two animals from the sham-operated group did not survive the recovery period. After an eight-week recovery phase to ensure systemic estrogen deprivation in the OVX group, all surviving animals were divided into four experimental groups: sham control diet (CD) group (n = 14; sham-operated females fed a CD), OVX CD group (n = 15; OVX females fed a CD), sham KD group (n = 14; sham-operated females fed a KD), and OVX KD group (n = 15; OVX females fed a KD). Both the CD and KD were formulated according to previously published compositions [55] by Top Class Enterprises LLP, New Delhi, India, using quality-matched macro and micronutrients. The nutrient composition of both diets is provided in the supplementary data file **(Table S1)**. To align feeding with the natural nocturnal activity of the animals, both diets were provided under a time-restricted schedule, with food access limited to 12h during the dark phase. Water was available *ad libitum* throughout the experimental period. Dietary interventions were continued for one month. Thereafter, neurobehavioral assessments were conducted by a blinded observer in a sound-attenuated room.

## Neurobehavioral studies

### Open field test

The open field test (OFT) was performed to assess locomotor activity, exploratory and anxiety-like behavior. Mice were acclimated to the behavioral testing room for 1h before the experiment. Each mouse was gently placed at the center of a square arena (55 × 55 × 55 cm) and allowed to explore for 10 min. Movements were recorded by an overhead camera and analyzed using ANY-maze™ software. After each trial, the arena was cleaned with 70% (v/v) ethanol and allowed to dry completely to eliminate residual scent cues. The number of entries into the central arena served as an index of anxiety-like behavior, while the total distance moved assessed locomotor activity.

### Elevated plus maze test

The elevated plus maze (EPM) test was conducted to assess anxiety-like behavior. Mice were acclimated to the behavioral testing room for 1h before the experiment. The apparatus consisted of a plus-shaped maze elevated 72.4 cm above the floor, containing two open arms (50.8 × 10.2 cm) and two closed arms (50.8 × 10.2 cm). Each mouse was gently placed at the center of the maze facing an open arm and allowed to explore freely for 10 min. Behavior was recorded using a video camera and analyzed with ANY-maze™ software. The apparatus was cleaned with 70% (v/v) ethanol and dried thoroughly between trials to eliminate residual scent cues. The time spent and the number of entries in the open and closed arms were measured to assess anxiety-like behavior, where time spent and the number of entries in open arms were inversely proportional to anxiety levels.

### Marble burying test

The marble burying test (MBT) was used to assess repetitive or compulsive-like behavior. Each mouse was placed individually in a cage measuring 29 × 17.5 cm containing 18 marbles evenly spaced on a 5 cm deep layer of bedding. A lid was secured on the cage to prevent escape. Mice were left undisturbed for 30 minutes, after which the number of marbles buried was counted. A marble was considered buried when at least two-thirds of its surface was covered by bedding. Higher numbers of buried marbles indicate increased compulsivity.

### Forced swimming test

The forced swimming test (FST) was used to assess depressive-like behavior. A transparent cylindrical tank (30 cm height, 10 cm diameter) was filled to three-quarters with water maintained at 23 ± 2°C. Each mouse was placed individually in the tank, and its activity was recorded for 6 minutes using ANY-maze™ software. The duration of immobility was measured and used as an indicator of depressive-like behavior.

### Nesting behavior

Nesting behavior was assessed to evaluate the self-care abilities of mice, following a previously described method [56] with minor modifications. Each mouse was housed individually for 36h in a cage containing 5g of cotton nesting material. The animals were left undisturbed during this period. After 36h, nests were examined independently by two observers and scored as follows: 0-no nest; 1-primitive flat nest, a pad-shaped structure slightly elevating the mouse above the bedding; 2-more elaborate nest involving warping and biting of the material; 3-well-formed cup-shaped nest with shredded material woven into walls; 4-hooded nest forming a hollow sphere with a single opening. The average score from both observers was used for further analysis.

### Blood glucose levels

Animals were subjected to a time-restricted feeding schedule in which food was removed during the light (rest) phase and provided during the dark (active) phase. Blood glucose levels were measured just before the onset of feeding at night, following the fasting period during the day. Blood samples were collected by tail prick, and glucose concentrations were determined using a handheld glucometer (Accu-Chek^®^ Active Blood Glucose Meter, Roche Diagnostics GmbH, Mannheim, Germany) from a small drop of blood.

### Blood and tissue collection

Blood samples were collected in a sterilised heparinized tube, and whole blood was left to clot at room temperature (RT) for 30 min before centrifugation at 3000 rpm for 15 min at 4°C. The clear plasma was separated and aliquoted into sterile microcentrifuge tubes, then stored at -80°C until further analysis. Likewise, brain and liver tissues intended for endpoint molecular analysis were rapidly dissected and stored at -80°C. A subset of samples for histopathological analysis was immersion-fixed in formalin, followed by cryoprotection using sequential 10% (w/v) and 30% (w/v) sucrose solutions.

### Estradiol levels

Plasma estradiol (E2) was quantified using an ELISA kit following the manufacturer’s instructions. Briefly, 50 µL of plasma and 50 µL of the biotinylated detection antibody were added to the pre-coated ELISA plate wells and incubated at 37°C. After incubation, the wells were washed three times with the supplied wash buffer (1 min each) and then incubated with 100 µL of HRP-conjugated antibody for 30 min at 37°C. The wells were washed five times, followed by the addition of 90 µL of the substrate reagent and 15 min incubation at 37°C in the dark. The reaction was stopped by adding 50 µL of stop solution, and absorbance was measured at 450 nm using a microplate reader. A standard curve prepared from the provided estradiol standards was used to determine the plasma estradiol levels.

### Corticosterone levels

Plasma corticosterone levels were measured using an ELISA kit according to the manufacturer’s protocol. Briefly, 50 µL of plasma and 50 µL of biotinylated antibody were added to the pre-coated ELISA plate wells and incubated at 37°C. After incubation, wells were washed three times with the supplied wash buffer (1 min each) and then incubated with 100 µL of HRP-conjugated antibody for 30 min at 37°C. Following five washes, 90 µL of substrate reagent was added, and plates were incubated in the dark at 37°C for 15 min. The reaction was stopped with 50 µL stop solution, and absorbance was recorded at 450 nm using a microplate reader. A standard curve generated from the kit’s corticosterone standards was used to calculate the plasma corticosterone levels.

### Ketone body estimation

β-hydroxybutyrate (BHB) levels were measured in plasma using a colorimetric assay kit following the manufacturer’s instructions. Briefly, 50 µL of plasma was mixed with 50 µL of the developer solution provided in the kit and incubated at 25°C in the dark for 30 min. Absorbance was recorded at 455 nm using a microplate reader. A standard curve was prepared using the supplied BHB standards to quantify BHB levels in plasma samples.

### Biochemical analysis

Plasma biochemical parameters were analyzed using the Cobas^®^ 8000 modular analyzer series automated analyzer, which utilizes photometric, electrochemiluminescence, and ion-selective electrode detection technologies to quantify analytes. Calibration curves were generated according to manufacturer instructions, and routine instrument maintenance and calibration logs were maintained. The following parameters were quantified: total cholesterol (TC), triglycerides (TG), high-density lipoprotein cholesterol (HDL-C), low-density lipoprotein cholesterol (LDL-C), aspartate aminotransferase (AST), alanine aminotransferase (ALT), alkaline phosphatase (ALP), total protein, albumin, urea, sodium, potassium, and chloride. Each assay required 150–200 µL of plasma. Samples were loaded into the analyzer, and automated readings for all parameters were recorded and tabulated. All assays were performed in duplicate to minimize technical variability, and any outlier values were re-assayed for confirmation.

### Cytokine array analysis

A membrane-based antibody array was used for the parallel determination of the relative levels of 111 mouse cytokines and chemokines. Plasma samples were processed following the manufacturer’s protocol. Array membranes were blocked with assay buffer 6 for 1h at RT, then incubated overnight at 4°C with plasma diluted in assay buffer 4. After incubation, membranes were washed three times with 1X wash buffer (10 min each) and incubated with the biotinylated detection antibody cocktail for 1h at RT. Following an additional washing step (three times, 10 min each), membranes were treated with streptavidin-HRP conjugate for 30 min at RT, washed again, and developed with the chemiluminescent reagents provided in the kit. Signal detection was performed using a chemiluminescence imaging system. Spot intensities were measured using ImageJ software after subtracting background values. Relative expression changes were calculated with respect to the sham CD group. Cytokines showing more than 1.5-fold upregulation or downregulation relative to the Sham CD group were considered for further analysis.

### Histopathology

Liver samples fixed in 10% neutral-buffered formalin were processed for histopathological evaluation of steatosis, fibrosis, and inflammation. Briefly, the tissues were washed overnight in running water, dehydrated through a graded ethanol series, cleared in xylene, and embedded in paraffin wax.

### Hematoxylin and eosin staining

Paraffin-embedded liver blocks were sectioned at 7 µm thickness, deparaffinized in xylene, and rehydrated through a graded ethanol series to distilled water. The sections were stained with Harris hematoxylin, rinsed in distilled water, and blued in lithium carbonate. Subsequently, the sections were counterstained with eosin, dehydrated through graded ethanol and xylene, and mounted using DPX mounting solution. Stained slides were examined under a bright-field microscope. Data represent the mean percentage of affected area by inflammation and steatosis across three sections per sample.

### Picrosirius red (PSR) staining

Paraffin-embedded liver sections (7 µm) were deparaffinized in xylene and rehydrated through a graded ethanol series to distilled water. The sections were then stained with Weigert’s iron hematoxylin, rinsed in running tap water, and subsequently stained with 0.1% (w/v) Sirius red solution in picric acid. After rinsing briefly in 0.5% (v/v) acetic acid, the sections were dehydrated through graded ethanol, cleared in xylene, and mounted with DPX mounting solution. Fibrotic collagen fibers were visualized under bright-field light microscopy. Data represent the mean percentage of PSR-positive staining in the central region across three sections per sample.

### Periodic acid–Schiff (PAS) staining

Paraffin-embedded sections (7 µm) were deparaffinized in xylene and rehydrated through a graded ethanol series to distilled water. The sections were oxidized in 0.5% (v/v) aqueous periodic acid, rinsed in distilled water, and incubated in Schiff reagent. After washing in running tap water, sections were counterstained with Harris hematoxylin, blued in lithium carbonate, and rinsed again in distilled water. Finally, the sections were dehydrated through graded ethanol, cleared in xylene, and mounted using DPX mounting solution. Data represent the mean percentage of PAS-positive staining in the central region across three sections per sample.

### Real-time PCR

Total RNA was extracted from liver tissues using TRIzol reagent following the manufacturer’s protocol. Complementary DNA (cDNA) was synthesized from 1 µg of total RNA using the RevertAid First Strand cDNA Synthesis Kit according to the recommended instructions. Gene-specific primers were designed using Primer3Plus software (supplementary data file, **Table S2**). Quantitative real-time PCR (qRT-PCR) was performed using SYBR Green Master Mix on a real-time PCR detection system. β-actin served as the internal control. Each reaction was run in technical duplicates to ensure reproducibility, and the mean value was used for analysis. Relative gene expression levels were calculated using the 2^^ΔΔCT^ method [57].

### DNA isolation from fecal samples and 16S rRNA gene sequencing

Genomic DNA was isolated from fecal samples using the kit, strictly adhering to the manufacturer’s protocol to ensure high yield and purity. The hypervariable V4 region of the 16S rRNA gene was amplified for sequencing using the established primer pair 341F (5’-CCTAYGGGRBGCASCAG-3’) and 806R (5’-GGACTACNNGGGTATCTAAT-3’). Libraries were prepared utilizing the DNA library prep kit, incorporating necessary adapters and indices. The resulting amplicon libraries were pooled and sequenced using 2 x 300bp paired-end reads on an Illumina MiSeq platform.

### 16S data analysis

Raw metagenomic reads obtained from Illumina sequencing were filtered to remove adapters and low-quality sequences using Fastp (Version 0.24.0) [58] with parameters --cut_right_mean_quality 20 and --length_required 100. Denoising, chimaera removal, and inference of Amplicon Sequence Variants (ASVs) were performed using the DADA2 pipeline within QIIME 2 (v2024.10.1) [59]. Taxonomic classification was carried out using a QIIME 2-compatible classifier pre-trained on the Genome Taxonomy Database (GTDB) [60]. Microbial diversity was analysed using alpha (within-sample) and beta (between-sample) diversities. Alpha diversity was analysed using the Mann-Whitney U test, whereas beta diversity was done using Principal Coordinate Analysis (PCoA) on Bray-Curtis distances, with statistical significance performed by PERMANOVA (999 permutations). Further, to identify differentially abundant taxa, Mann-Whitney U tests were performed on phylum-level abundances, using the Benjamini-Hochberg false discovery rate (FDR) correction (adjusted p<0.05). Diversity composition and its relative abundance were visulaized using bar plots and a heatmap of the top 25 taxa at the species level using Python (v3.11) with matplotlib, seaborn, and scikit-bio.

### RNA extraction from the medial prefrontal cortex and sequencing

Mice were transcardially perfused with cold PBS, and the medial prefrontal cortex was dissected on ice. Tissue was homogenized under RNase-free conditions, and total RNA was extracted using the RNeasy Mini Kit with optional on-column DNase treatment. RNA yield and purity were initially assessed by NanoDrop spectrophotometry. Library preparation and sequencing were performed by Eurofins Genomics, Bangalore, India. RNA integrity was further evaluated using the Agilent TapeStation with High Sensitivity RNA ScreenTape. QC-passed samples were used for paired-end library construction with the NEBNext^®^ Ultra™ II Directional RNA Library Prep Kit for Illumina. Briefly, mRNA was enriched using poly-T magnetic beads, enzymatically fragmented, and converted to first-strand cDNA, followed by second-strand synthesis to generate double-stranded cDNA. After purification with AMPure XP beads, dscDNA was A-tailed, adapter-ligated, and amplified by limited-cycle PCR. Final libraries were purified and assessed on the Agilent 4200 TapeStation using High Sensitivity D1000 ScreenTape for size distribution and concentration. All QC-qualified libraries were then subjected to cluster generation and high-throughput sequencing on the Illumina NovaSeq X Plus platform

### RNA-Seq data processing and analysis

Raw sequencing reads were initially subjected to quality assessment using FastQC v0.12.1. Adapter sequences and low-quality bases were trimmed using Trimmomatic v0.39 under standard quality filtering parameters, and post-trimming quality improvements were confirmed with MultiQC v1.30. High-quality reads were subsequently aligned to the mouse reference genome using the STAR aligner v2.7.11b. Alignment quality was assessed using SAMtools v1.22 and subcommand flagstat, qualimap v2.3 and summarized via MultiQC, showing consistently high mapping rates across all samples (∼99%), indicating excellent alignment success and high-quality sequencing data suitable for downstream differential expression analysis. Gene-level counts were obtained using featureCounts v2.1.1 with the corresponding GTF annotation file, yielding the primary raw gene count dataset. Transcript-level normalized expression profiles (FPKM and TPM) were estimated using StringTie v2.7.11b.

The raw read count matrix was further used for downstream differential expression analysis. Low-abundance genes were filtered to reduce background noise, followed by variance-stabilizing transformation (VST). DESeq2 v1.42.0 (R v4.4.3) was employed to identify differentially expressed genes (DEGs) using the Wald test, applying thresholds of |log₂ fold change| ≥ 1 and adjusted p-value < 0.05. Principal component analysis (PCA) was carried out on the VST-transformed dataset to assess overall sample clustering and group separation.

To interpret the biological significance of DEGs, Gene Ontology (GO) enrichment analyses were performed, highlighting modules related to neuronal, metabolic, molecular, neurobehavioral, immunological, and hormonal functions. These GO modules were visualized through heatmaps, Venn diagrams, volcano plots, network plots, and enrichment maps, while module-based average expression scores were also computed to capture pathway-level expression trends.

### Statistical analysis

Statistical analyses were performed using GraphPad Prism 8.1.2 (GraphPad Software, La Jolla, CA, USA). Data are presented as mean ± standard error of the mean (SEM). Two-way analysis of variance (ANOVA) was employed to evaluate the main effects of diet, hormone and diet X hormone interaction, followed by a Bonferroni post hoc test for multiple comparisons. For 16s RNA gene expression data analysis, alpha diversity was assessed using the Mann–Whitney U-test. Beta diversity was assessed using PERMANOVA (999 permutations) on Bray–Curtis distances. Differentially abundant phyla were analyzed using the Mann–Whitney U test with Benjamini–Hochberg False Discovery Rate (FDR) correction. Statistical significance was defined as a p ≤ 0.05.

## Compliance with Ethical Standards

## Statements and Declarations

### Funding

This work was supported by the Department of Biotechnology, Government of India, in the form of a research CORE grant to MK and fellowships to AKR and NG from the National Agri-Food and Biomanufacturing Institute (BRIC-NABI).

### Competing Interests

Authors have no competing financial interests.

### Data availability statement

The data supporting this article have been included as part of the Supplementary Information.

### Authors’ contributions

**AKR** performed the animal surgeries, behavioral assessments, real-time PCR, histopathological analysis, primary data analysis, and prepared the manuscript**. NG** analyzed the transcriptomic data and contributed to writing the method and results sections related to this data**. MMA** analyzed the gut microbiota data and assisted in writing the related sections**. PS** carried out the biochemical analyses. **AKR** provided supervision and analytical support for the gut microbiota experiments. **JK** supervised the acquisition of biochemical data. **MK** conceptualized the study, supervised the research, analyzed all data, wrote the manuscript, and finalized the manuscript.

### Ethics approval

Institutional Animal Ethics Committee (IAEC) of BRIC-NABI approved experimental protocol (NABI/2039/CPCSEA/IAEC/2023/06).

### Consent to participate

Not applicable

### Consent for publication

Not applicable

## Acknowledgements

The author thanks the Executive Director (BRIC-NABI) for the facilities. The authors are also thankful to the BRIC-NABI PARAMSMRITI Supercomputing facility for providing computing resources. Illustrations were created by using BioRender, an online scientific image illustration tool (BioRender.com).

## Notes

### Competing Interest Statement

The authors have declared no competing interest.

